# Auxin-salicylic acid seesaw regulates the age-dependent balance between plant growth and herbivore defense

**DOI:** 10.1101/2024.10.23.619823

**Authors:** Wen-Hao Han, Feng-Bin Zhang, Shun-Xia Ji, Kai-Lu Liang, Jun-Xia Wang, Xiao-Ping Fan, Shu-Sheng Liu, Xiao-Wei Wang

## Abstract

Plants must fine-tune their needs for growth and defense. According to the Plant Vigor Hypothesis, younger, more vigorous plants tend to be more susceptible to herbivores compared to older, mature plants, yet the molecular mechanisms underlying this dynamic remain elusive. Here, we uncover a hormonal crosstalk framework that orchestrates the age-related balance between plant growth and herbivore defense. We demonstrate that the accumulation of salicylic acid (SA), synthesized by *Nicotiana benthamiana* phenylalanine ammonia-lyase 6 (NbPAL6), dictates insect resistance in adult plants. The expression of *NbPAL6* is driven by the key transcription factor, NbMYB42, which is regulated by two interacting auxin response factors, NbARF18La/b. In juvenile plants, higher auxin levels activate *Nb*miR160c, a microRNA that simultaneously silences *NbARF18La/b*, subsequently reducing *NbMYB42* expression, lowering SA accumulation, and thus weakening herbivore defense. Furthermore, excessive SA accumulation in juvenile plants enhances defense but antagonizes auxin signaling, impairing early growth. Our findings suggest a seesaw-like model that balances growth and herbivore defense depending on the plant’s developmental stage.

## INTRODUCTION

Plants, being sessile organisms, must constantly navigate hostile environments to survive and reproduce, facing a wide range of abiotic and biotic stresses. Over time, they have evolved sophisticated systems to detect, respond to, and defend against environmental changes. However, these defenses come at a significant cost, often resulting in trade-offs with growth and development ^1,2^. Among the most damaging threats to plants are herbivorous insects, which can attack at any stage of plant development. While tissue-chewing insects cause direct damage by consuming plant tissues, phloem-feeding insects pierce the epidermis to delicately extract nutrient-rich phloem sap using needle-like mouthparts. In response to such threats, plants have evolved both inducible and constitutive defense mechanisms ^3,4^. Induced defenses are triggered in response to specific stimuli, while constitutive defenses involve continuous accumulation of insecticidal components throughout the plant’s growth and development ^5,6^. Given that insect attacks are often sporadic, plants must finely balance their investment in growth with the need for defense. However, how this balance is maintained at the molecular level remains poorly understood.

In the context of plant-herbivore interactions, the Plant Vigor Hypothesis (PVH) posits that herbivores preferentially target young, vigorous plants over older, mature ones ^7,8^. At the same time, plants tend to develop increased resistance to biological stresses, such as pathogens and herbivores, as they age—a phenomenon known as age-related resistance (ARR) ^9–13^. These observations suggest that plants may prioritize rapid growth early in life, tolerating some degree of herbivore damage, before shifting their focus to defense mechanisms later in development ^14^. However, the precise mechanisms underlying PVH and ARR in the context of herbivores remain largely unexplored. Given the dual necessity for plants to grow and defend themselves, a better understanding of how this balance is optimized holds significance to both ecological theory and agricultural production.

Phytohormones play a pivotal role in regulating various aspects of plant growth, development, and defense, often through intricate crosstalk mechanisms ^15,16^. Auxin, the first phytohormone discovered, along with other phytohormones, orchestrates plant growth and development while also modulating stress responses ^16^. Among stress-related phytohormones, jasmonic acid (JA), salicylic acid (SA), abscisic acid (ABA), and ethylene (ET) are key players, particularly in response to herbivory, with JA being central to resistance against tissue-chewing insects and SA being crucial for defense against phloem-feeding insects ^14,15^. The interaction between these phytohormones has complex effects on plants against insects ^15^. For instance, the JA-SA antagonistic relationship is a well-studied area, with each hormone modulating plant responses to different herbivores. Moreover, other phytohormones, including auxin, ET, ABA, gibberellins (GA), brassinosteroids (BR), and cytokinins (CK) are also involved in fine-tuning plant immunity by influencing JA and SA signaling pathways, especially in response to pathogens and abiotic stresses ^15^. The intricate mechanisms of phytohormonal interplay involve multiple layers of regulation, including transcriptional control by transcription factors (TFs), post-transcriptional regulation by small RNAs (sRNAs), and control of protein activity ^17–20^. Despite these advances, it remains unclear how plants integrate hormonal signals to balance growth and defense at different developmental stages.

The whitefly, *Bemisia tabaci* (Gennadius), is a species complex, several species of which are of agricultural importance worldwide ^21–24^. A few species of this whitefly complex are highly polyphagous, exhibiting remarkable adaptability to different hosts ^25,26^. Being phloem-feeders, whiteflies use their piercing-sucking mouthparts delicately to extract nutrients, making juvenile, vigorous plants particularly vulnerable to attacks and yield losses ^25,27,28^. Understanding the underlying mechanisms that govern plant growth-defense balance could be key to developing sustainable whitefly control strategies.

*Nicotiana benthamiana* has become an important model organism in plant molecular biology and biotechnology, serving as a bio-factory for vaccines and pharmacological compounds ^29,30^. Our previous studies have shown that *N. benthamiana* exhibits strong attraction to and lethal effects on several Hemiptera and Thysanoptera insects, including whiteflies, winged aphids, and thrips, highlighting its potential as a dead-end trap plant for field pest control ^31–33^. However, we observed that the insecticidal efficacy of *N. benthamiana* varied with the plant’s development stage. Upon further investigation, we found that beyond environmental influences, this age-related defense capacity was closely tied to the developmental stage, suggesting a complex regulatory mechanism involving ARR. These findings underscore the utility of *N. benthamiana* as a system to study ARR and provide a foundation for its application in pest management.

In this study, we demonstrate that auxin-SA crosstalk mediates the balance between plant growth and herbivore defense at different developmental stages. We reveal that the establishment of ARR in *N. benthamiana* and several other Solanaceae species is closely tied to SA levels. Using molecular biology and biochemical approaches, we elucidate that the NbARF18La/b-NbMYB42-NbPAL6 signaling module plays a critical role in the age-related regulation of SA accumulation. Furthermore, we find that the age-dependent auxin-miRNA160c module also contributes to this process. Our findings provide new insights into PVH and lay the theoretical groundwork for developing innovative pest control strategies against whiteflies.

## RESULTS

### Age-regulated resistance against phloem-feeding insects in Solanaceae plants

To compare the defense levels of plants at different ages against phloem-feeding insects, we conducted bioassays of *B. tabaci* whiteflies, *Myzus persicae* aphids, and *Frankliniella occidentalis* western flower thrips on several Solanaceae species, including *N. benthamiana*, *Solanum lycopersicum* (tomato), and *Nicotiana tabacum* (tobacco). In *N. benthamiana*, both the survival rate and fecundity of whiteflies were significantly lower on adult plants (35 days old, with 8–9 true leaves) compared to juvenile plants (21 days old, with 3–4 true leaves) (Figure 1A and 1B). Similarly, whiteflies performed better on juvenile tobacco (21 days old, 3–4 true leaves) and tomato (21 days old, 3–4 true leaves) plants compared to adult plants (35 days old, 6–7 true leaves for tobacco; 8–9 true leaves for tomato) (Figure 1A and 1B). Additionally, western flower thrips showed higher survival rates, and aphids produced more offspring on juvenile plants compared to adult plants (Figure 1C and 1D). These results demonstrate ARR against phloem-feeding insects in Solanaceae plants.

**Figure 1.**
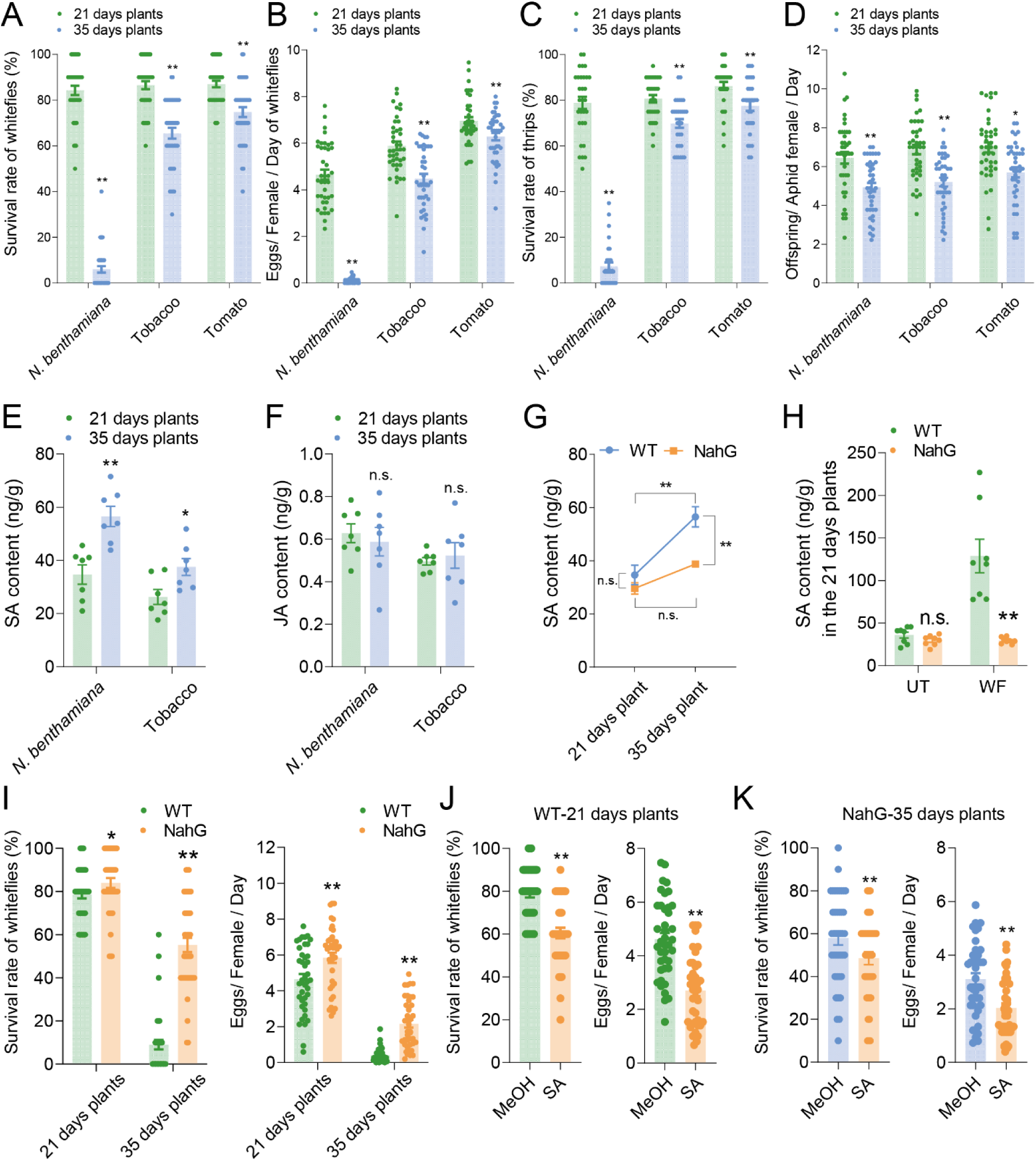
Plant age-related resistance against insects is regulated by SA. (A and B) Survival rate and fecundity of whiteflies on 21-day-old and 35-day-old *N. benthamiana*, tobacco, and tomato plants. (C) Survival rate of thrips on 21-day-old and 35-day-old *N. benthamiana*, tobacco, and tomato plants. (D) Fecundity of aphids on *N. benthamiana*, tobacco, and tomato plants at 21-day and 35-day ages. (E and F) SA and JA levels in *N. benthamiana* and tobacco at different ages. (G) Temporal dynamics of SA content in wild-type and NahG *N. benthamiana* plants. (H) SA levels in 21-day-old wild-type and NahG *N. benthamiana* plants upon whitefly attack. (I) Survival rate and fecundity of whiteflies on 21-day-old and 35-day-old wild-type and NahG *N. benthamiana* plants. (J) Survival rate and fecundity of whiteflies on SA-treated (1 mM) 21-day-old wild-type *N. benthamiana* plants. (K) Survival rate and fecundity of whiteflies on SA-treated (1 mM) 35-day-old NahG *N. benthamiana* plants. Values are means ± SE, n = 40 for A, B, C, D, I, J, and K, n = 7 for E, F, and G, n = 8 for H. Student’s *t*-test (two-tailed) was used for significant difference analysis. n.s., not significant. \**P* < 0.05, \*\**P* < 0.01.

### SA-dependent ARR against phloem-feeding insects

To explore the mechanism underlying ARR, we focused on *N. benthamiana*, which exhibited the most pronounced age-dependent insect resistance. We examined whether the defense phytohormones SA and JA were involved in ARR by comparing the basal levels of SA and JA in 21-day-old and 35-day-old *N. benthamiana* plants. SA levels were higher in adult plants, while JA levels showed no significant differences between juvenile and adult plants (Figure 1E and 1F). This trend was also observed in tobacco (Figure 1E and 1F). Additionally, the expression of *pathogenesis-related 1* (*NbPR1*), a downstream marker of SA signaling, was significantly upregulated in adult plants, indicating enhanced SA signaling (Figure S1A).

To further investigate the role of SA in ARR, we used *N. benthamiana* transgenic plants expressing the *NahG* gene, which encodes *salicylate hydroxylase* from *Pseudomonas putida*, an enzyme that depletes SA accumulation ^34^. Unlike wild-type plants, NahG plants were unable to accumulate sufficient SA as they aged (Figure 1G), and the mRNA levels of *NbPR1* were also significantly reduced in 35-day-old plants (Figure S1B). Moreover, whitefly infestation failed to induce SA accumulation in NahG plants (Figure 1H). Bioassays revealed that whiteflies performed better on both juvenile and adult NahG plants compared to wild-type plants (Figure 1I), indicating that both whitefly-inducible and growth-accumulated SA are critical for plant defense against whiteflies. Exogenous SA application further confirmed SA’s role in ARR. Treatment with SA increased *NbPR1* expression and enhanced whitefly defense in 21-day-old wild-type plants (Figure 1J and Figure S1C), as well as in 35-day-old NahG plants (Figure 1K). These results collectively confirm that ARR in *N. benthamiana* against whiteflies is SA-dependent.

### SA-mediated ARR depends on the expression of the SA synthesis gene NbPAL6

In plants, SA is synthesized via two distinct pathways: the phenylalanine ammonia lyase (PAL) pathway and the isochorismate synthase (ICS) pathway ^35^ (Figure 2A). To assess whether gene expression in these pathways changes with plant age, we performed reverse transcription quantitative PCR (RT-qPCR). The results showed no significant differences in the transcript levels of *NbICS1* and *NbICS2*, or the the two TFs, *CaM-Binding Protein 60g* (*CBP60g*) and *Systemic Acquired Resistance Deficient 1* (*SARD1*), which activate the ICS pathway (Figure 2B). Similarly, components of the ICS pathway such as *Enhanced Disease Susceptibility 5* (*EDS5*) and *AvrPphB Susceptible 3* (*PBS3*) were downregulated with plant age, while *Enhanced Pseudomonas Susceptibility 1* (*EPS1*) remained unchanged (Figure 2B). Given these findings, we shifted our focus to the PAL pathway.

**Figure 2.**
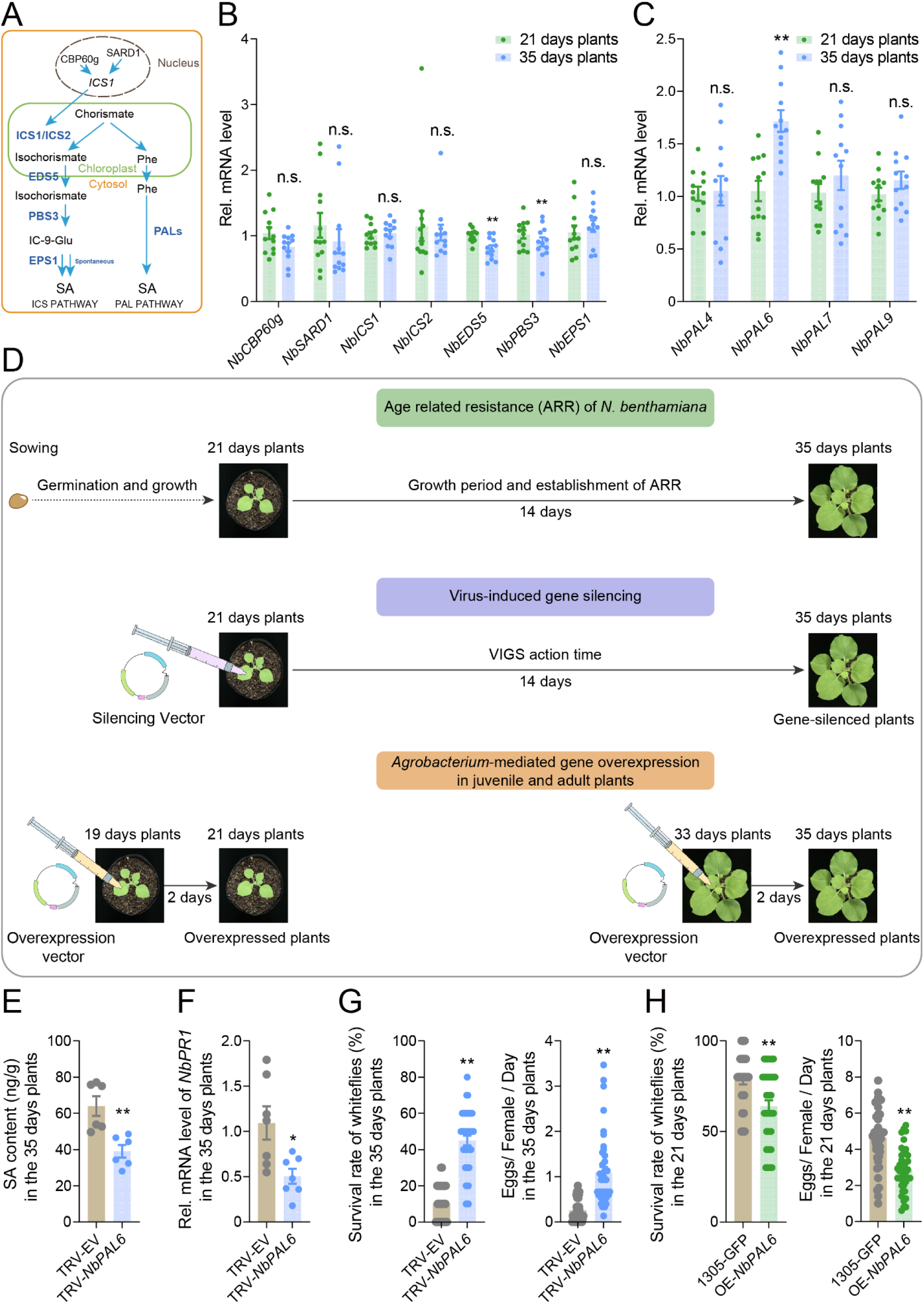
ARR depends on the expression of the SA synthetic gene *NbPAL6*. (A) SA synthesis pathways in plants. (B) Expression levels of genes in the ICS pathways in 21-day-old and 35-day-old *N. benthamiana* plants. (C) Expression levels of *NbPAL* genes in 21-day-old and 35-day-old *N. benthamiana* plants. (D) Experimental design of VIGS and gene overexpression in *N. benthamiana* plants. In the VIGS system, the pTRV2 vector is injected into 21-day-old plants, activating the silencing vector to inhibit target gene expression. Gene silencing typically became evident 14 days post-injection, aligning with the timeline for ARR establishment in *N. benthamiana*, making VIGS an effective tool for studying ARR-related gene functions. In comparison to control plants (receiving only the empty vector), 35-day-old plants subjected to gene silencing show reduced accumulation of target gene transcripts after reaching 14 days. Additionally, *Agrobacterium*-mediated gene overexpression enhances target gene mRNA levels in both 21-day-old and 35-day-old plants, allowing for gene function assessment across different developmental stages. For instance, when the pCAMBIA1305-GFP vector containing the target gene is introduced at 19 days, mRNA levels increase in the treated 21-day-old plants within two days. By comparing overexpressed plants to controls (receiving only the empty vector), the impact of target genes on resistance in 21-day-old plants can be assessed. This method was also used to evaluate the role of gene overexpression in enhancing resistance in 35-day-old plants. (E) SA levels in *NbPAL6*-silenced 35-day-old *N. benthamiana* plants. (F) Expression level of *NbPR1* in *NbPAL6*-silenced 35-day-old *N. benthamiana* plants. (G) Survival rate and fecundity of whiteflies on *NbPAL6*-silenced 35-day-old *N. benthamiana* plants. (H) Survival rate and fecundity of whiteflies on *NbPAL6*-overexpressed 21-day-old *N. benthamiana* plants. Values are mean ± SE, n = 12 for B and C; n = 7 for F; n = 6 for E; n = 40 for G and H. Student’s *t*-test (two-tailed) was used for significant difference analysis. n.s., not significant. \**P* < 0.05, \*\**P* < 0.01.

A search of the *N. benthamiana* genome database (https://www.nbenth.com/) identified nine *PAL* genes (*NbPAL1*-*NbPAL9*), which we classified into four phylogenetic branches, likely representing alleles from the allotetraploid ancestry of *N. benthamiana* (Figure S2 and Table S1). From these, we selected four *PAL* genes (*NbPAL4*, *NbPAL6*, *NbPAL7*, and *NbPAL9*) from the four branches respectively for further analysis. Expression profiling across developmental stages revealed that only *NbPAL6* showed increased transcription with plant age, while the other genes remained unchanged (Figure 2C).

To determine whether *NbPAL6* is involved in SA-mediated ARR, we employed virus-induced gene silencing (VIGS) and *Agrobacterium*-mediated overexpression to manipulate *NbPAL6* expression in *N. benthamiana* ^30,36^. The timing of these treatments was carefully designed to align with plant age requirements (Figure 2D).

In the VIGS system, we injected the pTRV2 vector into 21-day-old plants., which led to effective gene silencing within 14 days (Figure 2D). This period of transcriptional inhibition coincides with the establishment of ARR, making VIGS an ideal system for studying gene-related ARR (Figure 2D).

Compared to control plants (injected with an empty vector), 35-day-old VIGS-treated plants showed a significant reduction in target gene transcript accumulation after reaching 21 days of age. Subsequent validation experiments on the 35-day VIGS-treated plants provide valuable insights into the role of target genes in ARR establishment. Meanwhile, *Agrobacterium*-mediated overexpression can be used to enhance the mRNA level of target genes in both 21-day-old and 35-day-old plants, enabling examination of gene function in plants of different ages (Figure 2D). We injected 19-day-old plants with the pCAMBIA1305-GFP vector containing the target gene. Two days later, during the vector’s active period, elevated target gene mRNA levels were observed in the 21-day-old plant. By comparing overexpressed plants with control plants (injected with an empty vector), we could evaluate the effect of target genes on resistance in 21-day-old plants (Figure 2D). Similarly, this method can also verify the effect of target gene overexpression in 35-day-old plants (Figure 2D).

Our results showed that 35-day-old VIGS-*NbPAL6* plants could not accumulate SA to the same levels as control (VIGS-EV) plants, nor could they upregulate *NbPR1* expression (Figure 2E and 2F; Figure S3A). Moreover, these plants exhibited reduced resistance to whiteflies (Figure 2G). Conversely, overexpression of *NbPAL6* at 21 days of age significantly increased the *NbPR1* levels and improved plant defense against whiteflies (Figure 2H; Figure S3B and S3C). These findings indicate that the differential expression of *NbPAL6* between juvenile and adult stages regulates SA level and SA-mediated ARR against whiteflies in *N. benthamiana*.

### Age-regulated NbMYB42 activates NbPAL6 transcription

To identify the upstream TFs regulating *NbPAL6* expression, we screened the *NbPAL6* promoter (*NbPAL6pro*) using a yeast one-hybrid (Y1H) system with a cDNA library from *N. benthamiana*. NbMYB42, a TF belonging to the R2R3-MYB family, was identified. R2R3-MYB is known to regulate the phenylpropanoid metabolic pathway ^37–39^. Yeast transformants containing the plasmids pAbAi-*NbPAL6pro* and pGADT7-NbMYB42 were able to grow on SD/-Leu medium with 150 ng/ml AbA within 3 days, while control transformants with pAbAi-*NbPAL6pro* and pGADT7 did not (Figure 3A). This indicates that NbMYB42 can bind to the *NbPAL6* promoter. NbMYB42 was also found to localize in the nucleus (Figure 3B), suggesting its role as a nuclear TF. To assess its role in activating *NbPAL6* expression in vivo, we performed the GUS staining assays. The results showed that *NbPAL6pro::GUS* co-infiltrated with *35S::NbMYB42* exhibited significantly higher GUS staining levels compared to the empty vector control in *N. benthamiana* leaves (Figure 3C). Similarly, dual-luciferase reporter assays demonstrated increased luciferase activity when *NbPAL6pro::LUC* was co-expressed with *35S::NbMYB42* (Figure 3D). These results confirmed that NbMYB42 activates *NbPAL6* promoter expression.

**Figure 3.**
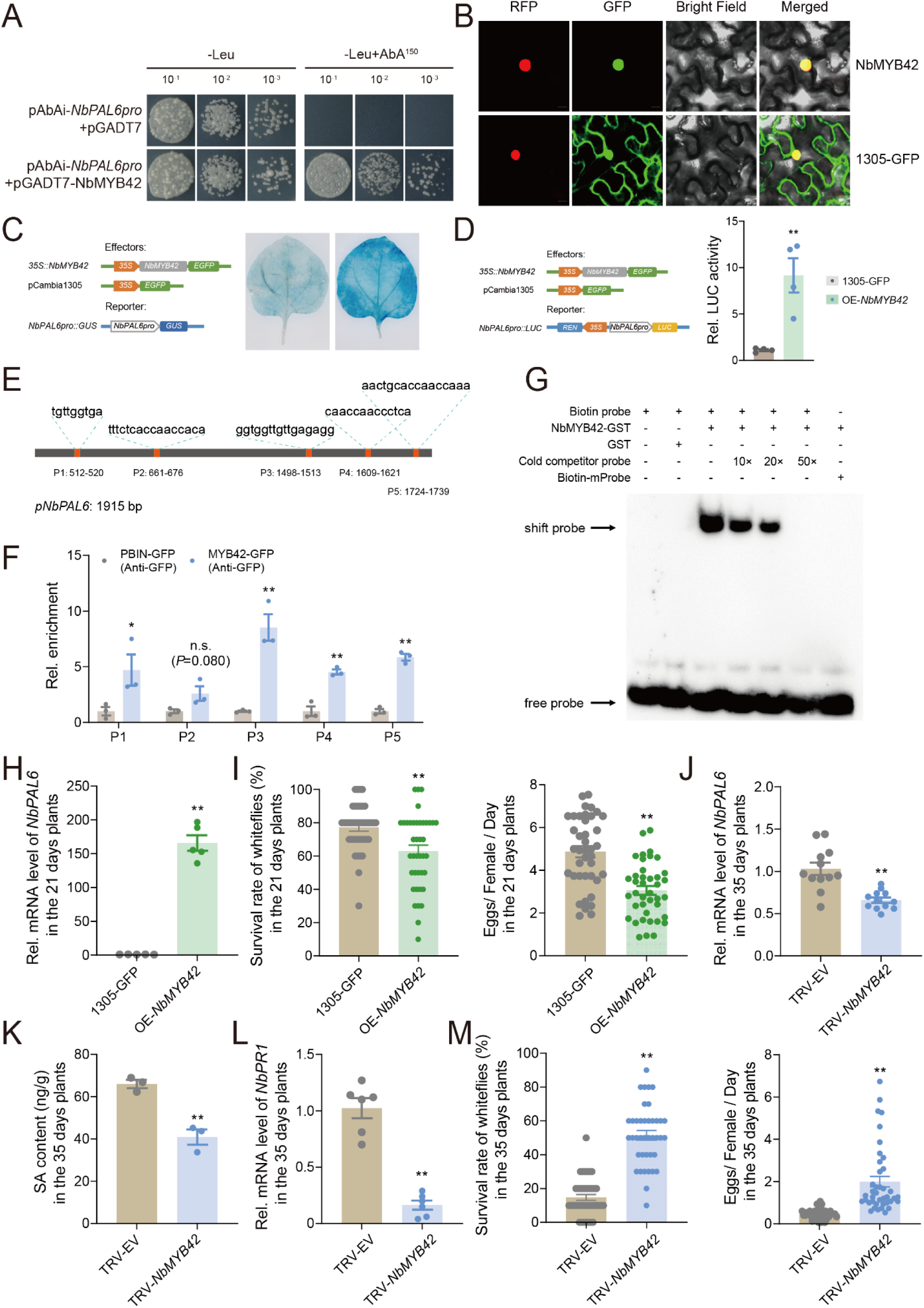
NbMYB42 directly binds to the *NbPAL6* promoter to activate its transcription. (A) Yeast one-hybrid assay of the interaction between the *NbPAL6* promoter and NbMYB42. Gold yeast cells co-transformed with pAbAi-*NbPAL6pro* and pGADT7-NbMYB42 plasmids were cultured on SD/-Leu medium with or without 150 ng/ml AbA for 3 days. The empty vector pGADT7 served as a negative control. (B) Subcellular localization of NbMYB42 in *N. benthamiana*. NbMYB42-GFP (green fluorescent protein) was transiently expressed in *N. benthamiana* line H2B-RFP, with red fluorescent protein (RFP) as a nucleus marker and the empty *35S::GFP* vector used as a control. Scale bar: 20 μm. (C) Analysis of the interaction between the *NbPAL6* promoter and NbMYB42 in *N. benthamiana* leaves using a GUS staining assay. Diagrams on the left illustrate the reporter and effector vectors: *35S*, CaMV *35S* promoter; *EGFP*, *enhanced green fluorescent protein*; *GUS*, β*-glucuronidase*. Representative photographs from several replicates are shown. (D) Analysis of the interaction between the *NbPAL6* promoter and NbMYB42 in *N. benthamiana* leaves using a dual-luciferase reporter assay. Diagrams on the left illustrate the reporter and effector vectors. *REN*, *Renilla luciferase*; *LUC*, *firefly luciferase*. The activity of REN was used as an internal control. (E) Schematic of the *NbPAL6* promoter, indicating possible binding sites P1 to P5 for NbMYB42. (F) ChIP-qPCR assay of the enrichment of NbMYB42-GFP at the *NbPAL6* promoter with an anti-GFP antibody. PCR primer pairs were designed based on the position shown in (E). Element P3 (GGTGGTTGTTGAGAGG) is significantly enriched in NbMYB42-GFP plants. (G) EMSA results show NbMYB42-GST binding to the *NbPAL6* promoter in vitro. The NbMYB42-GST protein was incubated with a biotin-labeled probe to assess binding capacity. Labeled probes were incubated with and without GST protein as negative controls, and 10-, 20-, and 50-fold excess unlabeled probes were used as cold competitors. Biotin-mProbe was the labeled probe containing mutations in the P3 element of the *NbPAL6* promoter. (H) Expression level of *NbPAL6* in *NbMYB42*-overexpressed 21-day-old *N. benthamiana* plants. (I) Survival rate and fecundity of whiteflies on *NbMYB42*-overexpressed 21-day-old *N. benthamiana* plants. (J) Expression level of *NbPAL6* in *NbMYB42*-silenced 35-day-old *N. benthamiana* plants. (K) SA levels in *NbMYB42*-silenced 35-day-old *N. benthamiana* plants. (L) Expression level of SA downstream gene *NbPR1* in *NbMYB42*-silenced 35-day-old *N. benthamiana* plants. (M) Survival rate and fecundity of whiteflies on *NbMYB42*-silenced 35-day-old *N. benthamiana* plants. Values are mean ± SE, n = 4 for D and F; n = 5 for H; n = 40 for I and M; n = 12 for J; n = 3 for K; n = 6 for L. Student’s *t*-test (two-tailed) was used for significant difference analysis. n.s., not significant. \**P* < 0.05, \*\**P* < 0.01.

Next, we explored whether NbMYB42 directly binds to specific sites within the *NbPAL6pro*. Using the Jasper database (https://jaspar.elixir.no/), we identified five potential MYB binding sites (P1–P5) (Figure 3E). To validate binding in vivo, we conducted chromatin immunoprecipitation (ChIP)-qPCR using plants overexpressing *NbMYB42* with GFP tag. We designed primers specific to the five binding sites to assess their interaction with NbMYB42 (Table S2). The experiments were carried out using an anti-GFP antibody. Binding activity was observed at P1, P3, P4, and P5, but not P2, in the immunoprecipitated chromatin, indicating a significant accumulation of NbMYB42-GFP at these sites compared to the GFP control (Figure 3F). Furthermore, electrophoretic mobility shift assay (EMSA) with recombinant NbMYB42-GST, purified from *Escherichia coli*, confirmed the binding activity in vitro. The MYB element site P3, which showed the strongest binding ability in the ChIP-qPCR test, was selected to prepare the biotin-labeled probe according to its sequence (Table S2). We observed that the NbMYB42-GST protein was able to bind directly to the P3 probe, while the mutated probe eliminated NbMYB42-GST binding (Figure 3G).

Additionally, an unlabeled P3 probe was prepared as the cold competitors. With the increase of competitor probes (10, 20, and 50 ×), the shifted band corresponding to the P3-probe and the NbMYB42 complex gradually disappeared (Figure 3G), suggesting the binding capacity was greatly reduced by the addition of unlabeled cold competition probe. These results collectively show that NbMYB42 specifically binds to the MYB *cis*-element on the *NbPAL6* promoter.

We then studied whether NbMYB42 influences ARR against whiteflies by regulating NbPAL6-mediated SA accumulation. Overexpressing *NbMYB42* in 21-day-old *N. benthamiana* plants increased *NbPAL6* expression more than 150-fold compared to control plants (Figure 3H and Figure S3D). The SA-downstream *NbPR1* was also significantly upregulated (Figure S3E). This overexpression enhanced whitefly resistance in 21-day-old plants (Figure 3I). Conversely, silencing *NbMYB42* in 35-day-old plants suppressed *NbPAL6* expression (Figure 3J and Figure S3F), resulting in reduced SA and *NbPR1* levels (Figure 3K and 3L). Notably, the *NbMYB42*-silenced 35-day-old *N. benthamiana* plant showed significantly reduced whitefly defense compared to the control (Figure 3M). These results indicate that NbMYB42 plays a crucial role in ARR in *N. benthamiana* by regulating *NbPAL6* expression and SA accumulation.

### Two NbARFs activate the expression of NbMYB42

We observed that *NbMYB42* expression was significantly higher in 35-day-old plants compared to 21-day-old plants (Figure 4A), prompting further investigation into how *NbMYB42* expression is regulated during *N. benthamiana* development. Using a Y1H assay, we screened for TFs that regulate *NbMYB42* expression and identified two auxin response factors (ARFs) with distinct evolutionary relationships, which we named NbARF18La and NbARF18Lb (Figure S4A). Yeast strains harboring the pAbAi-*NbMYB42pro* and pGADT7-NbARF18La or pGADT7-NbARF18Lb constructs were able to grow on SD/- Leu medium supplemented 150 ng/ml AbA, whereas yeast strains containing pAbAi-*NbMYB42pro* and a control pGADT7 plasmid could not (Figure 4B), suggesting that both NbARF18La and NbARF18Lb bind to the *NbMYB42* promoter.

**Figure 4.**
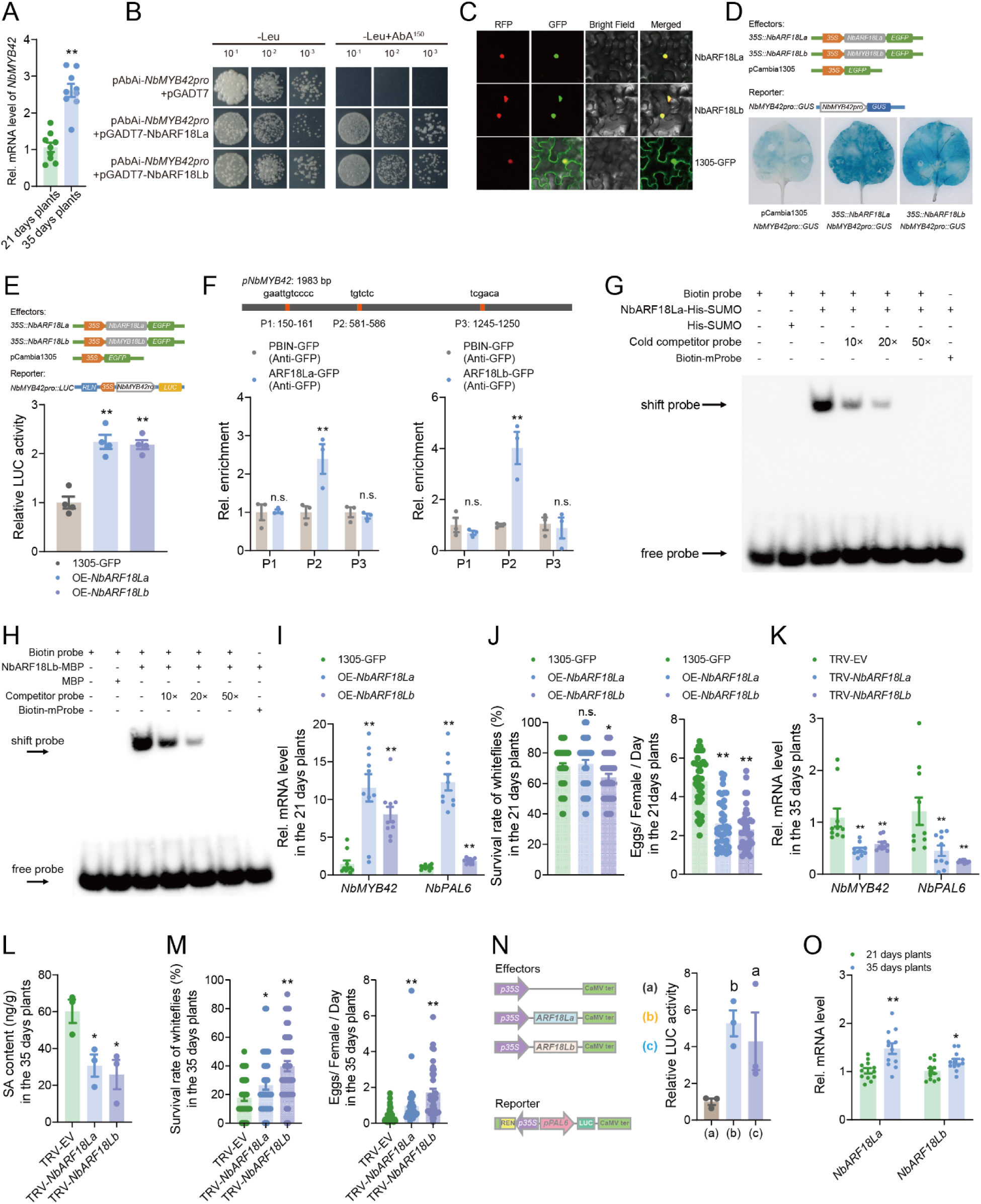
NbARF18La and NbARF18Lb activate *NbMYB42* expression. (A) Expression levels of *NbMYB42* in 21-day-old and 35-day-old *N. benthamiana* plants. (B) Yeast one-hybrid assay of the interaction between the *NbMYB42* promoter and NbARF18La/b. Y2HGold yeast cells co-transformed with pAbAi-*NbMYB42pro* and pGADT7-NbARF18La/b plasmids were cultured on SD/-Leu medium with or without 150 ng/ml AbA for 3 days. The empty vector pGADT7 was used as a negative control. (C) Subcellular localization of NbARF18La and NbARF18Lb in *N. benthamiana*. NbARF18La/b-GFP fusion proteins were transiently expressed in *N. benthamiana* line H2B-RFP. Red fluorescent protein (RFP) served as a nucleus marker, and the empty *35S::GFP* vector was used as a control. Scale bar: 20 μm. (D) GUS staining assay analyzing the interaction between the *NbMYB42* promoter and NbARF18La/b in *N. benthamiana* leaves. Diagrams on the top show the reporter and effector vectors. (E) Dual-luciferase reporter assay analyzing the interaction between the *NbMYB42* promoter and NbARF18La/b in *N. benthamiana* leaves. REN activity was used as an internal control. (F) Top: Schematic of the *NbMYB42* promoter showing the distribution of possible NbARF binding sites. Bottom: ChIP-qPCR assay of the enrichment for NbARF18La/b-GFP protein at the *NbMYB42* promoter with an anti-GFP antibody. The PCR primer pair is based on the positions shown above. Element P2 (TGTCTC) is significantly enriched in NbARF18La/b-GFP plants. (G and H) EMSA showed the binding of NbARF18La-His-SUMO and NbARF18Lb-MBP to the *NbMYB42* promoter in vitro. Proteins were incubated with a biotin-labeled fragment of the *NbMYB42* promoter (P2, TGTCTC). The labeled probe was incubated with and without His-SUMO or MBP protein as negative controls and 10-, 20- and 50-fold excess unlabeled probes were used as cold competitor probes. Biotin-mProbe refers to the labeled probe with mutations. (I) Expression levels of *NbMYB42* and *NbPAL6* in *NbARF18La/b*-overexpressed 21-day-old *N. benthamiana* plants. (J) Survival rate and fecundity of whiteflies on *NbMYB18a/b*-overexpressed 21-day-old *N. benthamiana* plants. (K) Expression levels of *NbMYB42* and *NbPAL6* in *NbARF18La/b*-silenced 35-day-old *N. benthamiana* plants. (L) SA levels in *NbARF18La/b*-silenced 35-day-old *N. benthamiana* plants. (M)Survival rate and fecundity of whiteflies on the *NbARF18La/b*-silenced 35-day-old *N. benthamiana* plants. (N) LUC activity was driven by the *NbPAL6* promoter when co-expressed with different effectors. (O) Expression levels of *NbARF18La/b* in 21-day-old and 35-day-old *N. benthamiana* plants. Values are mean ± SE, n = 9 for A; n = 4 for E; n = 3 for F, L and N; n = 10 for I and K; n = 40 for J and M; n = 12 for O. Student’s *t*-test (two-tailed) was used for significant difference analysis. n.s., not significant. \**P* < 0.05, \*\**P* < 0.01.

Phylogenetic analysis showed that NbARF18La and NbARF18Lb are closely related to ARF16 from *Arabidopsis thaliana* (Figure S4A). Amino acid sequence analysis revealed conserved B3-type DNA-binding domains in the N-termini of both NbARF18La and NbARF18Lb and dimerization domains in their C- termini ^40–43^ (Figure S4B). Subcellular localization analysis confirmed that both NbARFs are localized in the nucleus (Figure 4C), indicating their function as nuclear TFs. To verify the activation of *NbMYB42* by these NbARFs, we conducted GUS staining and dual-luciferase reporter assays. Co-expression of *35S::NbARF18La* and *35S::NbARF18Lb* with the reporter *NbMYB42pro::GUS* in *N. benthamiana* leaves significantly increased *GUS* expression (Figure 4D), and dual-luciferase reporter assay showed a significant up-regulation in LUC/REN ratio when *35S::NbARF18La* and *35S::NbARF18Lb* were co-expressed with the *NbMYB42pro::LUC* reporter (Figure 4E). These results collectively demonstrate that NbARF18La and NbARF18Lb activate *NbMYB42* expression.

### DNA-binding specificity of NbARF18La and NbARF18Lb

To explore how the two NbARFs specifically bind to the *NbMYB42* promoter, we screened for potential auxin response elements (AuxRE) using the JASPAR database and identified three regions (P1, P2, and P3) in the promoter (Figure 4F). We designed primers specific to these regions (Table S2) and performed ChIP-qPCR using GFP-tagged NbARF18La/b and anti-GFP antibodies. The results revealed a significant enrichment of the P2 region, but not P1 and P3, in the immunoprecipitated chromatin, indicating that NbARF18La/b bind to the P2 AuxRE region (TGTCTC) in the *NbMYB42* promoter (Figure 4F).

To confirm the binding specificity, we conducted an EMSA assay. The recombinant NbARF18La-His-SUMO fusion protein bound to the P2 probe of the *NbMYB42* promoter, while no binding occurred with a mutant probe (TGTCTC to AAAAAA) (Figure 4G). Additionally, competition assays using unlabeled probes verified the specificity of binding (Figure 4G). Similarly, the NbARF18Lb-MBP fusion protein bound to the same promoter region (P2), but not to the mutant probe, and competition with unlabeled probes also reduced the binding strength (Figure 4H). Together, these results indicate that NbARF18La and NbARF18Lb specifically bind to the same AuxRE site in the *NbMYB42* promoter.

### NbARF18La and NbARF18Lb interaction and self-association

Previous studies have shown that ARF TFs can form complexes through self-interactions or by interacting with other TFs ^40–42,44^. To explore potential interactions between NbARF18La and NbARF18Lb or self-interactions of either of the TFs, we performed yeast two-hybrid (Y2H) assays. Yeast cells expressing NbARF18La-AD/NbARF18Lb-BD grew on selective selective media (SD/-Leu-Trp-His-Ade), indicating that NbARF18La and NbARF18Lb can interact with each other (Figure S5A). Additionally, NbARF18La was able to interact with itself in Y2H assays (Figure S5A).

We then confirmed these interactions in vivo using a bimolecular fluorescence complementation (BiFC) assay. GFP fluorescence was detected in *N. benthamiana* leaves co-expressing *NbARF18La* and *NbARF18Lb* or *NbARF18La* alone (Figure S5B). In vitro pull-down assays further validated these interactions, as NbARF18La-His exhibited an affinity for NbARF18Lb-MBP, but no interaction was observed in the negative control (Figure S5C). These results demonstrate that NbARF18La and NbARF18Lb physically interact and NbARF18La can self-interact, suggesting that these interactions may be important for cooperatively regulating *NbMYB42* expression.

### NbARFs regulate plant resistance to whiteflies in an age-related manner

Next, we examined the role of NbARF18La and NbARF18Lb in ARR to whiteflies in *N. benthamiana*. Overexpression of both *NbARFs* significantly upregulated expression of *NbMYB42*, *NbPAL6*, and the downstream *NbPR1* gene in 21-days-old plants (Figure 4I; Figure S6A to S6C), which led to enhanced resistance to whiteflies (Figure 4J). In contrast, silencing *NbARF* genes via VIGS in 35-day-old *N. benthamiana* plants resulted in substantially reduced expression of *NbMYB42* and *NbPAL6* (Figure 4K; Figure S6D and S6E), as well as significantly decreased endogenous SA levels and reduced *NbPR1* expression (Figure 4L; Figure S6F). Correspondingly, silencing *NbARFs* impaired the defense response against whiteflies in adult plants (Figure 4M).

To further investigate how NbARFs regulate *NbPAL6* expression, we co-expressed *NbARF18La/b* with a LUC reporter fused to the *NbPAL6* promoter, which significantly enhanced LUC activity (Figure 4N), indicating that NbARF18La/b regulate *NbPAL6* by modulating the expression of *NbMYB42*. Furthermore, expression of *NbARF18La/b* was significantly higher in 35-day-old plants than in 21-day-old plants (Figure 4O), indicating that the NbARF18La/b-NbMYB42-NbPAL6 module is regulated in an age-dependent manner.

### Auxin suppresses plant defense by repressing SA accumulation in juvenile plants

Given the findings above, we questioned why SA-mediated defense is not fully established during the juvenile stage of plants. Since defense-related ARF TFs are key components downstream of the auxin response, we postulated that auxin is involved in this process. To test this hypothesis, we first measured the levels of indole-3-acetic acid (IAA) in 21-day-old and 35-day-old plants, and found that IAA levels were significantly higher in juvenile plants (Figure 5A). In parallel, we observed elevated expression levels of the auxin synthesis genes, *TRYPTOPHAN AMINOTRANSFERASE RELATED 2* (*NbTAR2*) and *YUCCA 8* (*NbYUC8*), as well as the auxin-responsive gene *NbIAA29*, in juvenile plants (Figure 5B).

**Figure 5.**
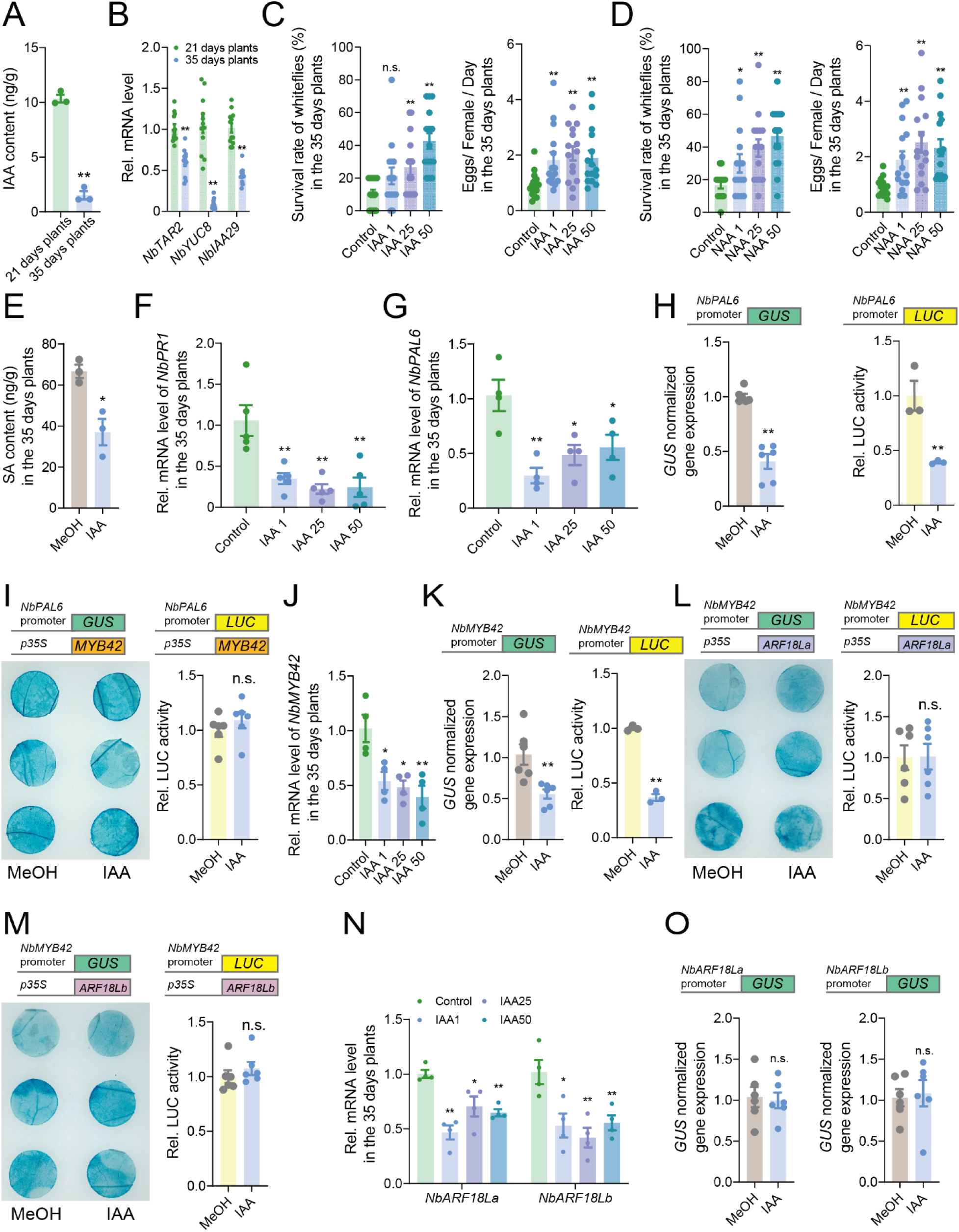
Auxin suppresses NbARF18La/b-MYB42-mediated SA accumulation. (A) IAA levels in 21-day-old and 35-day-old *N. benthamiana* plants. (B) Expression level of *NbTAR2*, *NbYUC8*, and *NbIAA29* in 21-day-old and 35-day-old *N. benthamiana* plants. (C, D) Survival rate and fecundity of whiteflies on IAA/NAA-treated (1, 25, and 50 μM) 35-day-old *N. benthamiana* plants. (E) SA levels in IAA-treated (25 μM) 35-day-old *N. benthamiana* plants. (F) Expression level of *NbPR1* in IAA-treated (1, 25, and 50 μM) 35-day-old *N. benthamiana* plants. (G) Expression level of *NbPAL6* in IAA-treated (1, 25, and 50 μM) 35-day-old *N. benthamiana* plants. (H) Top: Schematic of the *GUS* reporter gene. Bottom: *GUS* reporter gene expression in mock- and IAA-treated (25 μM) *pNbPAL6::GUS N. benthamiana* plants. (I) Top: Schematic of the *GUS* and *LUC* reporter genes and effector *NbMYB42*. Bottom: GUS / LUC activity co-expressed with *NbMYB42* in mock- or IAA-treated (25 μM) *pNbPAL6::GUS N. benthamiana* plants. (J) Expression level of *NbMYB42* in IAA-treated (1, 25, and 50 μM) 35-day-old *N. benthamiana* plants. (K) Top: Schematic of the *GUS* reporter genes. Bottom: *GUS* reporter gene expression in mock- and IAA-treated (25 μM) *pNbMYB42::GUS N. benthamiana* plants. (L) Top: Schematic of the *GUS* and *LUC* reporter gene and effector *NbARF18La*. Bottom, GUS / LUC activity co-expressed with *NbARF18La* in mock- and IAA-treated (25 μM) *pNbMYB42::GUS* and *pNbMYB42::LUC N. benthamiana* plants. (M) Top: Schematic of the *GUS* and *LUC* reporter gene and effector *NbARF18Lb*. Bottom: GUS / LUC activity co-expressed with *NbARF18Lb* in mock- and IAA-treated (25 μM) *pNbMYB42::GUS* and *pNbMYB42::LUC N. benthamiana* plants. (N) Expression level of *NbARF18La/b* in IAA-treated (1, 25, and 50 μM) 35-day-old *N. benthamiana* plants. (O) Top: Schematic of the *GUS* reporter gene. Bottom: *GUS* reporter gene expression in mock- and IAA-treated (25 μM) *pNbARF18La/b::GUS N. benthamiana* plants. Values are mean ± SE, n = 3 for A and E; n = 12 for B; n = 15 for C and D; n = 5 for F; n = 4 for G, J and N; n = 6 for H, K, and O (GUS normalized gene expression), n = 3 for H and K (Relative LUC activity); n = 6 for I, L, and M (Relative LUC activity). Student’s *t*-test (two-tailed) was used for significant difference analysis. n.s., not significant. \**P* < 0.05, \*\**P* < 0.01.

To assess the effect of auxin on plant defense, we treated 35-day-old plants, which are known for their high resistance to whiteflies, with natural IAA and the auxin-like plant growth regulator, 1-naphthalene acetic acid (NAA). Bioassays revealed that the whiteflies performed better on IAA- and NAA-treated plants, indicating that auxin negatively impacts plant defense (Figure 5C and 5D). Furthermore, IAA treatment significantly suppressed both the SA levels (Figure 5E) and the expression of the SA-responsive gene *NbPR1* (Figure 5F) in the 35-day-old plants.

To further clarify the role of auxin in repressing SA-based resistance, we treated 21-day-old plants with two auxin biosynthesis inhibitors: _L_-kynurenine (_L_-Kyn), a competitive substrate inhibitor, and Yucasin (5- (4-chlorophenyl)-4H-1,2,4-triazole-3-thiol), an effective IAA biosynthesis inhibitor ^45^. Plants treated with these auxin inhibitors showed increased resistance to whiteflies (Figure S7A and S7B), as well as higher expression levels of *NbPR1* (Figure S7C). These results confirm that auxin suppresses SA biosynthesis and downstream signaling pathways in juvenile *N. benthamiana* plants, thereby repressing SA-mediated defense against whiteflies.

### Auxin suppresses NbPAL6 and NbMYB42 expression by repressing NbARF levels

To explore how auxin suppresses SA accumulation, we focused on the age-regulated SA synthesis gene *NbPAL6*. Auxin treatment significantly reduced *NbPAL6* expression in 35-day-old plants (Figure 5G), whereas treatment with an auxin inhibitor enhanced *NbPAL6* expression in 21-day-old plants (Figure S7D). To investigate the underlying mechanism, we used *NbPAL6* promoter-driven *GUS* and *LUC* reporter, which showed decreased transcript levels upon auxin treatment, indicating that auxin suppresses *NbPAL6* expression through transcriptional repression (Figure 5H).

We hypothesized that auxin may inhibit *NbPAL6* transcription by suppressing the activity of NbMYB42. However, *NbPAL6* promoter-linked *GUS* and *LUC* reporter assays in IAA-treated plants revealed no significant impact on reporter expression (Figure 5I), suggesting that auxin does not affect the transactivation ability of NbMYB42.

Next, we assessed whether auxin affects *NbMYB42* mRNA levels. RT-qPCR revealed a significant reduction in *NbMYB42* mRNA levels in auxin-treated 35-day-old plants (Figure 5J), whereas auxin inhibitor treatment enhanced *NbMYB42* expression in 21-day-old plants (Figure S7E). A reporter-GUS assay further confirmed that auxin represses *NbMYB42* transcription (Figure 5K). These results indicate that auxin represses *NbMYB42* transcription, thereby regulating downstream *NbPAL6* expression.

To understand how auxin represses *NbMYB42* transcription, we examined upstream TFs, NbARF18La and NbARF18Lb. Surprisingly, auxin treatment did not affect the activation of *NbMYB42* by NbARF18La/b (Figure 5L and 5M). However, auxin significantly decreased the transcript levels of *NbARF18La/b* in 35- day-old plants (Figure 5N), with auxin inhibitors increasing their expression in 21-day-old plants (Figure S7F). These results indicate that auxin downregulates NbARF18La/b, which in turn suppresses *NbMYB42*. Further, we examined whether auxin repressed *NbARFs* through transcriptional inhibition.

Importantly, using promoter-*GUS* reporters, we found that auxin did not reduce *NbARF18La/b* mRNA levels through transcriptional repression (Figure 5O). Therefore, it appears that auxin suppresses *NbARF* mRNA levels through a post-transcriptional regulatory mechanism.

### Auxin-inducible miRNA NbmiR160c targets NbARF mRNA

We explored the mechanisms by which auxin suppresses the transcript levels of *NbARF18La* and *NbARF18Lb* via post-transcriptional regulation. Previous studies have shown that plant miRNAs can negatively regulate the transcripts of *ARF* genes ^46,47^, and that auxin treatment can enhance miRNA accumulation in plants ^48^. Therefore, we hypothesized that auxin-inducible, age-related miRNAs may be responsible for the post-transcriptional regulation of these *ARF* genes. To test this hypothesis, we analyzed miRNA sequencing data from 21-day-old *N. benthamiana* plants and, through target prediction analysis, identified a miRNA, *Nb*miR160c, which potentially targets the transcripts of *NbARF18La* and *NbARF18Lb*. Target site analysis revealed that *Nb*miR160c specifically binds to the coding regions of *NbARF18La* (1572 bp to 1590 bp) and *NbARF18Lb* (2099 bp to 2107 bp) (Figure 6A and 6B). To validate this interaction, we artificially overexpressed *Nb*miR160c in 35-day-old plants. This overexpression significantly reduced the mRNA levels of *NbARF18La* and *NbARF18Lb* (Figure 6C; Figure S8A and S8B), suggesting direct regulation of these transcripts by *Nb*miR160c.

**Figure 6.**
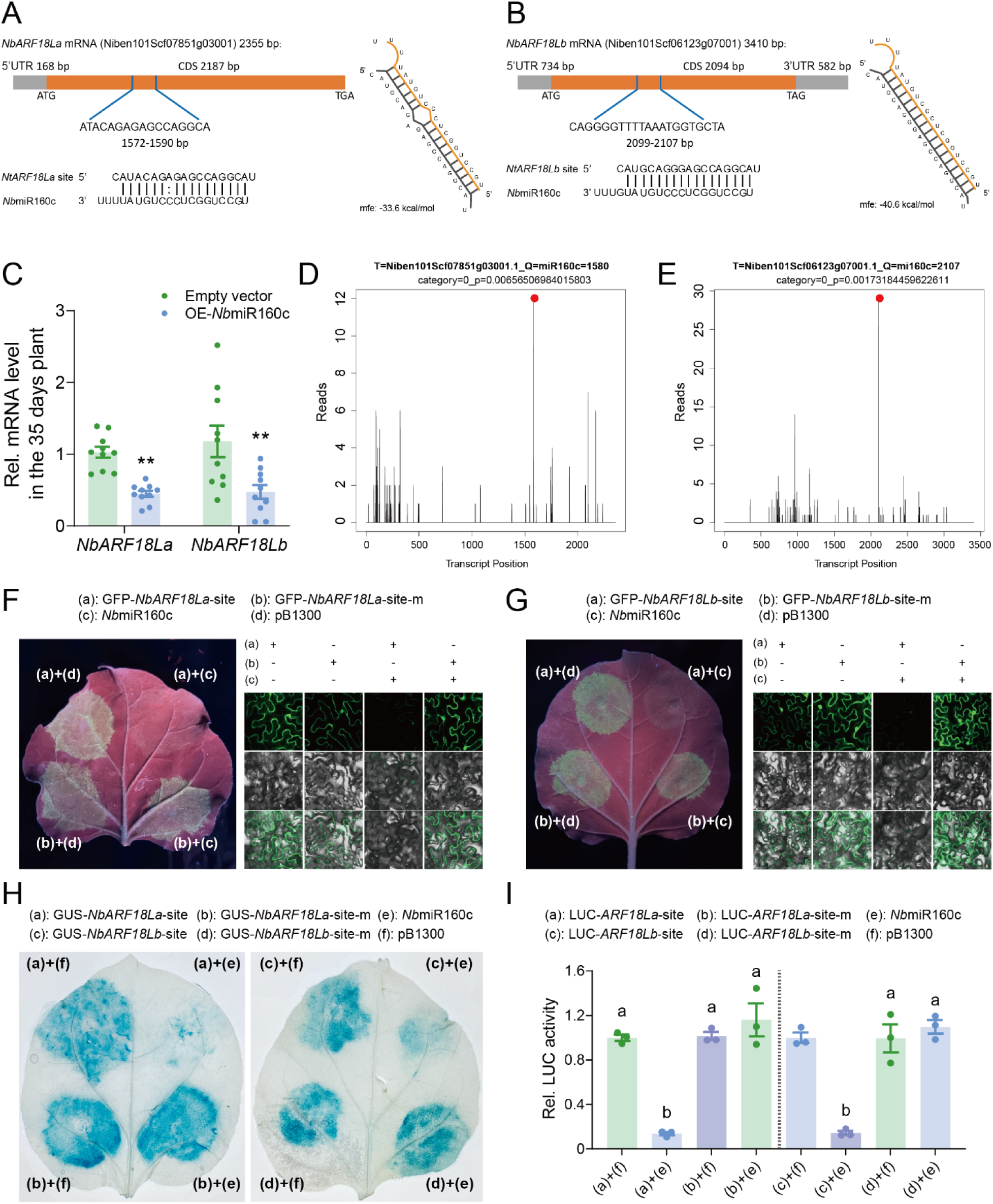
Auxin-induced microRNA160c targets *NbARF18La/b* transcripts. (A) *Nb*miR160c targets *N. benthamiana NbARF18La* mRNA. CDS: Coding sequence; UTR: Untranslated regions; ATG: Initiation codon; TAG and TGA: Termination codon. (B) *Nb*miR160c targets *N. benthamiana NbARF18Lb* mRNA. (C) Expression level of *NbARF18La/b* in *Nb*miR160c-expressing 35- day-old *N. benthamiana* plants. (D) Target plots (t-plots) of *Nb*miR160c and its target *NbARF18La* mRNA, validated via degradome sequencing. The red dot indicates the cleavage site of the *NbARF18La* transcript at 1572 bp, as analyzed by GSTAr (1.0) cleavage site prediction software. (E) T-plots of *Nb*miR160c and its target *NbARF18Lb* mRNA, showing cleavage at 2099 bp. (F, G) Camera (Left) and confocal (Right) images of GFP-NbARF18La/b target site and GFP-NbARF18La/b mutant site expression in *N. benthamiana* when *Nb*miR160c was overexpressed. Top: GFP; Middle, Bright-field; Bottom: GFP/bright-field overlay. Scale bars, 40 μm. (H) GUS phenotype observed by histochemical staining showing GFP-NbARF18La/b target site and GFP-NbARF18La/b mutant site expression in *N. benthamiana* with overexpressed *Nb*miR160c. (I) LUC activity displaying LUC-NbARF18La/b target site and LUC-NbARF18La/b mutant site expression. Values are mean ± SE, n = 10 for C; n = 3 for I. Student’s *t*-test (two-tailed) was used for significant difference analysis in C. \*\**P* < 0.01. One-way ANOVA followed by Fisher’s least significant difference (LSD) test was used for significant difference analysis in I. Bars with different lowercase letters indicate significant differences between treatments at *P* < 0.05.

Next, to pinpoint the exact cleavage sites of *NbARF18La* and *NbARF18Lb* by *Nb*miR160c in juvenile plants, we conducted degradome sequencing on samples from 21-day-old plants. The analysis confirmed that *Nb*miR160c cleaves *NbARF18La* and *NbARF18Lb* transcripts at the predicted target sites, as shown by the accumulation of cleavage products at these regions (Figure 6D and 6E). The red dots in Figure 6D and 6E highlight the cleavage sites, aligning with the *Nb*miR160c target predictions.

To further confirm the regulatory role of *Nb*miR160c, we cloned the target sites of *NbARF18La* and *NbARF18Lb* and fused them with a GFP tag. When these constructs were transiently expressed in *N. benthamiana* leaves, co-expression with *Nb*miR160c resulted in the suppression of GFP fluorescence (Figure 6F and 6G; Figure S8C and S8D). However, when the *Nb*miR160c target sites were mutated, *GFP* expression was unaffected, even in the presence of *Nb*miR160c (Figure 6F and 6G; Figure S8C and S8D). Additionally, in co-expression assays with GUS-tagged target sites, *Nb*miR160c suppressed the expression of wild-type GUS-tagged *NbARF18La* and *NbARF18Lb* target sites, while the mutant target sites were unaffected (Figure 6H). We observed similar results using LUC-tagged reporter constructs, where *Nb*miR160c suppressed the activity of the LUC reporter fused to the wild-type target sites of *NbARF18La* and *NbARF18Lb*, but not the mutant constructs (Figure 6I). Collectively, these results confirm that *Nb*miR160c directly targets the coding regions of *NbARF18La* and *NbARF18Lb*, leading to their post-transcription repression. This auxin-inducable miRNA thereby plays a crucial role in modulating *ARF* expression in *N. benthamiana*, particularly during the juvenile stage.

### NbmiR160c regulates the expression of NbMYB42 and NbPAL6

To explore the role of *Nb*miR160c in regulating the NbARF-NbMYB42 module-mediated SA defense against whiteflies, we first overexpressed *Nb*miR160c in 35-day-old plants. The overexpression resulted in a significant reduction in the mRNA levels of *NbMYB42* and *NbPAL6* (Figure 7A), as well as in SA content and downstream *NbPR1* expression (Figure 7B and Figure S8E). Notably, the *Nb*miR160c- expressing 35-day-old plants exhibited reduced defense against whiteflies (Figure 7C), indicating that *Nb*miR160c negatively regulates plant anti-insect defense.

**Figure 7.**
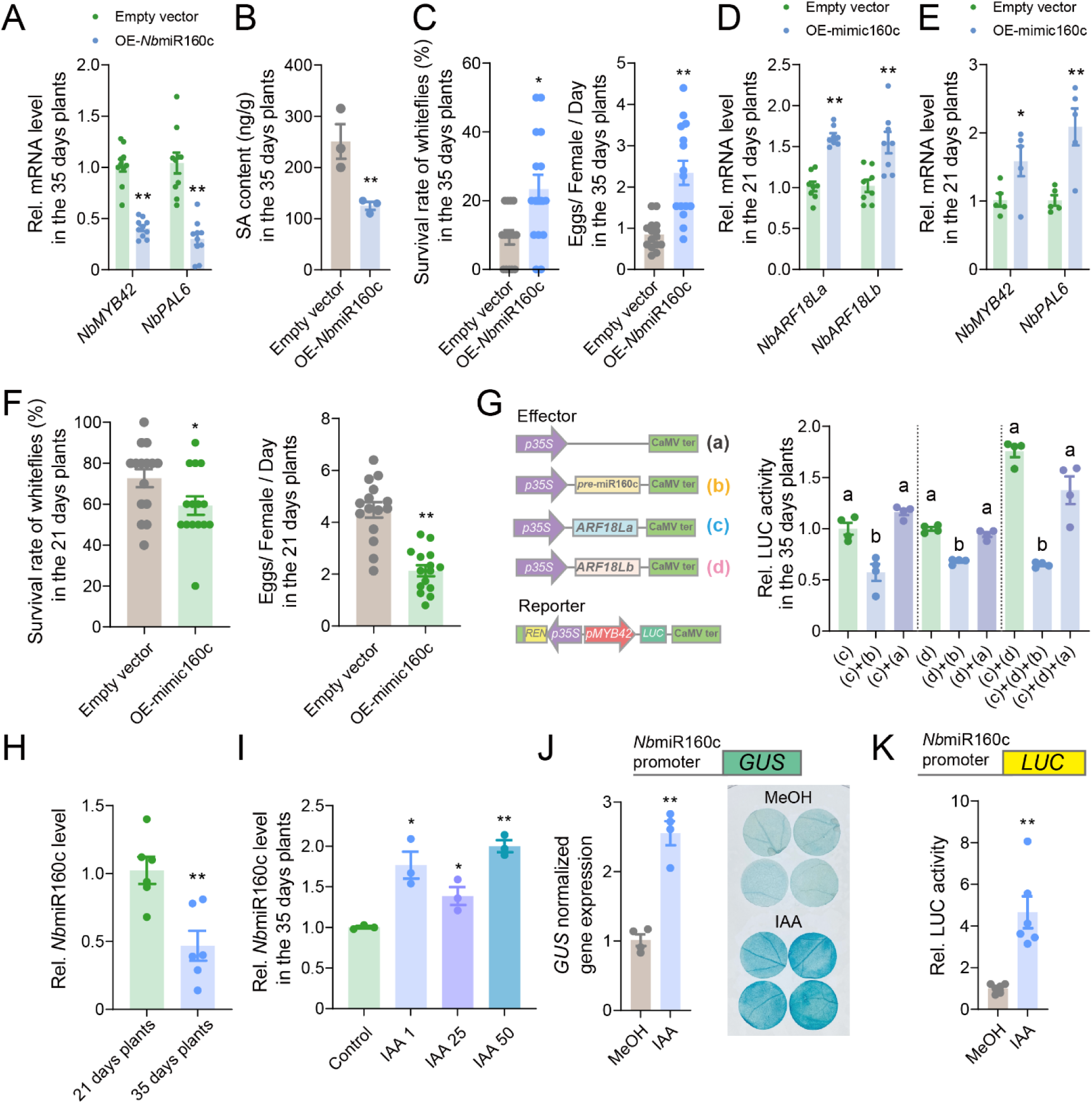
Auxin-induced microRNA160c negatively regulates plant defense. (A) Expression level of *NbMYB42* and *NbPAL6* in *Nb*miR160c-expressing 35-day-old *N. benthamiana* plants. (B) SA levels in *Nb*miR160c-expressing 35-day-old *N. benthamiana* plants. (C) Survival rate and fecundity of whiteflies on *Nb*miR160c-expressing 35-day-old *N. benthamiana* plants. (D) Expression level of *NbARF18La/b* in mimic160c-expressing 21-day-old *N. benthamiana* plants. (E) Expression level of *NbMYB42* and *NbPAL6* in mimic160c-expressing 21-day-old *N. benthamiana* plants. (F) Survival rate and fecundity of whiteflies on the mimic160c-expressing 21-day-old *N. benthamiana* plant. (G) Left: Schematic diagram showing effector and reporter vectors used in transient transcriptional activity assays in *N. benthamiana* leaves. Right: The LUC activity driven by the *NbMYB42* promoter when co-expressed with different effectors. (H) Expression level of *Nb*miR160c in 21-day-old and 35-day-old *N. benthamiana* plants. (I) *Nb*miR160c levels in IAA-treated (1, 25, and 50 μM) 35-day-old plants. (J) Top: Schematic of the *GUS* reporter gene; Bottom: *GUS* reporter gene expression and activity in mock- and IAA-treated (25 μM) *pNb*miR160c*::GUS N. benthamiana* plants. (K) Top: Schematic of the *LUC* reporter gene; Bottom: *LUC* reporter gene expression and activity in mock- and IAA-treated (25 μM) *pNb*miR160c*::LUC N. benthamiana* plants. Values are mean ± SE; n = 10 for A; n = 3 for B and I; n = 15 for C and F; n = 8 for D; n = 5 for E; n = 6 for H and K; n = 4 for G and J. Student’s *t*-test (two-tailed) was used for significant difference analysis in A, B, C, D, E, F, H, I, J, and K. \**P* < 0.05, \*\**P* < 0.01. One-way ANOVA followed by Fisher’s least significant difference (LSD) test was used for significant difference analysis in G. Bars with different lowercase letters indicate significant differences between treatments at *P* < 0.05.

To validate the role of *Nb*miR160c in repressing *NbARF18La/b* and downstream SA regulatory pathways in juvenile plants, we overexpressed an artificial target mimic of *Nb*miR160c (mimic160c) in 21- day-old plants to inhibit its function (Figure S8F). As expected, *NbARF18La* and *NbARF18Lb* transcript levels were significantly upregulated in the treated plants (Figure 7D). Moreover, the *NbMYB42* and *NbPAL6* mRNA levels (Figure 7E), SA levels (Figure S8G), and *NbPR1* expression were also significantly elevated in the mimic160c-overexpressing plants (Figure S8H). Correspondingly, bioassays showed that whiteflies performed poorly on the mimic160c-overexpressing plants, indicating enhanced defense (Figure 7F).

We then examined the effect of *Nb*miR160c on NbARF-promoted *NbMYB42* transcription in plants at juvenile and adult stages. In 35-day-old plants, overexpression of *Nb*miR160c repressed the activity of the *NbMYB42pro*-*LUC* reporter (Figure S9A), whereas overexpression of mimic160c enhanced *NbMYB42pro*-*LUC* reporter expression in 21-day-old plants (Figure S9B). When *Nb*miR160c was co-expressed with *NbARF18La*, *NbARF18Lb*, or both, *NbMYB42pro* reporter expression was significantly inhibited in 35-day-old plants (Figure 7G). These results demonstrate that *Nb*miR160c represses NbARF- activated *NbMYB42* transcription. Similarly, we confirmed that *Nb*miR160c is involved in similarly repressing *NbPAL6* transcription (Figure S9C to S9E). Collectively, these findings show that *Nb*miR160c negatively regulates *NbARF18La* and *NbARF18Lb*, thereby affecting the transcription of *NbMYB42* and its downstream target *NbPAL6*.

### Auxin promotes the NbmiR160c level by enhancing its expression

Next, we investigated the relationship between the *Nb*miR160c expression, plant age, and phytohormones. We observed that *Nb*miR160c levels were significantly higher in 21-day-old plants compared to 35-day-old plants (Figure 7H). Given that auxin content is typically higher in juvenile plants, we speculated that auxin might promote *Nb*miR160c expression. To test this hypothesis, we treated 35- day-old plants, which had naturally lower *Nb*miR160c levels, with IAA. The treatment significantly increased *Nb*miR160c levels (Figure 7I). Additionally, we cloned the promoter of the *Nb*miR160c precursor from the *N. benthamiana* genome and fused it to *GUS* and *LUC* reporters. Upon IAA treatment, we observed enhanced reporter expression (Figure 7J and 7K), confirming that auxin activates *Nb*miR160c transcription. These results indicate that auxin promotes *Nb*miR160c expression, thereby repressing the NbARF-NbMYB42-NbPAL6 module, which mediates whitefly defense in juvenile *N. benthamiana* plants.

### Excessive SA impacts early plant growth by antagonizing auxin

We further examined the physiological implications of auxin-mediated suppression of SA in juvenile plants. To assess the impact of excessive SA on early plant growth, we treated 21-day-old plants with exogenous SA. These treated plants showed stunted growth (Figure S10A) and, by the time they reached 35 days of age, had significantly reduced weight and height compared to untreated plants (Figure S10B and S10C). These results, along with our earlier findings, suggest that while excessive SA enhances whitefly defense in juvenile plants, it also negatively impacts plant early growth.

Since auxin and SA pathways are known to antagonize each other, we examined whether SA inhibited early growth by interfering with the auxin pathway. Our results show that SA treatment significantly reduced the expression of the auxin synthesis genes *NbTAR2* and *NbYUC8* (Figure S10D), lowered IAA levels (Figure S10E), and decreased downstream *NbIAA29* expression (Figure S10F) in juvenile plants. Conversely, the NahG plants, which have reduced SA levels, showed enhanced expression of auxin synthesis genes compared to wild-type plants at 35 days (Figure S10G). These results demonstrate that excessive SA in the early stages of plant development inhibits auxin accumulation, thereby hindering normal growth.

## DISCUSSION

In nature, plants exhibit enhanced defense against herbivorous insects as they age, with juvenile plants experiencing more frequent attacks than their adult counterparts. This observation has led to the development of the concepts of ARR and PVH; however, the precise mechanisms underlying these phenomena remain largely unknown. In this study, we elucidate the fine regulation of SA-mediated ARR against phloem-feeding whiteflies in plants (Figure 1). We highlight the importance of the PAL synthesis pathway in age-related intrinsic SA accumulation, identifying the key synthetic gene *NbPAL6* in *N. benthamiana* (Figure 2). Additionally, we identify the age-related upstream TFs NbMYB42 and NbARF18La/b that are involved in SA regulation (Figure 3 and Figure 4).

Furthermore, we provide insights into why SA-mediated resistance is not fully established in juvenile plants. Specifically, high levels of auxin in juvenile plants activate *Nb*miR160c, which silences the *NbARF18La/b* genes, ultimately impairing SA accumulation. This mechanism also prevents excessive SA from disrupting early plant growth. Our findings deepen the understanding of the PVH, demonstrating that juvenile plants exhibit lower herbivore tolerance while prioritizing urgent growth needs (Figure S11).

Our previous research has established the significant role of inducible SA in tobacco defense against whiteflies ^49^. In the current study, we found that the increased intrinsic SA levels associated with age also correlate with enhanced defense against whiteflies in adult plants (Figure 1). Although JA is crucial for plant defense against whiteflies ^49^, our findings indicate that the intrinsic JA levels did not vary with age across several Solanaceae species (Figure 1F), indicating that JA may not be a key phytohormonal signal in plant ARR against whiteflies. Recently, the role of SA in plant defense against phloem-feeding insects has gained attention ^50–52^, and our findings broaden the understanding of SA-mediated defenses from a novel perspective.

Moreover, we found the PAL pathway, rather than the ICS pathway, in SA biosynthesis changes at the transcriptional level with age in *N. benthamiana* (Figure 2B and 2C). Previous studies have shown the key role of *PAL* genes in SA accumulation in tobacco and other plant species ^38,53,54^. Here, we systematically analyzed *PAL* genes in *N. benthamiana* (Figure S2) and determined the age-related upregulation of *NbPAL6* expression (Figure 2C) and its role in age-related SA accumulation (Figure 2E). It is worth noting that our conclusion does not imply that *NbPAL6* is the sole *NbPAL* gene responsible for SA synthesis in *N. benthamiana*, as other PALs may contribute redundantly to SA synthesis. We suspect that additional NbPALs may participate in SA accumulation under various conditions, including biotic or abiotic stresses, warranting further investigation.

While extensive attention has been given to the upstream regulation of SA synthesis pathways, most studies have focused on the ICS pathway ^55^. The PAL pathway is known to contribute to the synthesis of numerous metabolic compounds in plants, such as flavonoids and lignin ^56^, but the regulation of *PAL* gene expression related to SA synthesis has been underexplored. For instance, in rice, the brown planthopper induces OsMYB30, which activates *OsPAL* gene expression, leading to increased SA and lignin accumulation and enhanced resistance ^38^. In our study, we demonstrated that the TF NbMYB42 activates *NbPAL6* expression by binding to its promoter, thereby enhancing SA levels and whitefly defense in an age-related constitutive, but not inducible, manner (Figure 3). Interestingly, in our previously published work, NbMYB42 was also shown to regulate lignin synthesis pathways in *N. benthamiana*, contributing to enhanced whitefly resistance ^39^. These observations combined highlight the necessity of MYB-PAL modules in plant SA and lignin synthesis, as well as their defense against phloem-feeding insects. In summary, our findings elucidate the mechanisms of age-related SA synthesis and regulation in plants, providing a valuable reference for future studies on the significance of age-enhanced SA in plant physiology and its induction in plant-pathogen and plant-insect interactions.

It is well established that SA and auxin signaling pathways often interact antagonistically ^15^. Elevated auxin levels correlate with reduced *PR* gene expression ^57,58^ and increased susceptibility to pathogens ^58–60^. In this study, we found that increased auxin levels can also impact plant defense against phloem-feeding insects by antagonizing SA levels (Figure 5A to 5F). Previous studies have indicated that auxin signaling may affect SA biosynthesis, although the underlying mechanisms remain largely unknown ^15,58^.

Our study revealed that auxin inhibits the expression of the key synthesis gene *NbPAL* and its transcriptional regulators, NbMYB42 and NbARF18La/b, thereby reducing SA biosynthesis (Figure 5G to 5O). In addition, repression of auxin signaling by SA has been demonstrated in studies on plant growth. For example, SA treatment decreased biomass in Arabidopsis, an effect that was abolished in the *auxin resistant 3* (*axr3*) mutant background, which harbors an auxin-negative regulator ^61^. Similarly, we found that increased SA levels influenced auxin accumulation and related gene expression (Figure S10D to S10F), leading to reduced growth in *N. benthamiana* plants (Figure S10A to S10C). Collectively, these findings deepen our understanding of the crosstalk between SA and auxin, particularly regarding upstream biosynthesis.

Studies have shown that ARFs activate MYB TFs involved in various essential physiological processes ^62–65^. Our study complementarily reports that the two ARFs, NbARF18La and NbARF18Lb, positively regulate SA biosynthesis by promoting SA-related MYB TFs, thereby enhancing plant defense against herbivores (Figure 4). Notably, the two ARFs we identified can interact with one another (Figure S5). Our findings indicate that auxin inhibits SA accumulation, positioning NbARFs as central hubs in this interaction. While ARFs are known participants in the antagonism between auxin and other phytohormones, such as ABA and JA ^43,66^, little is reported regarding their relationship with SA. Previous studies have shown that the auxin-induced ARF7 represses the expression of SA-downstream *PR* genes that regulate bacterial infections in Arabidopsis plants ^67^. In contrast, we found that in *N. benthamiana* auxin represses the transcriptional activators *NbARF18La/b* (Figure 6), thereby negatively affecting upstream SA signaling.

Additionally, this work reveals the mechanism by which auxin represses *NbARFs* through miR160c- mediated post-transcriptional regulation (Figure 6). Prior studies have demonstrated that several miRNAs target *ARF* transcripts to regulate growth and development. For example, in Arabidopsis, miR160 targets *ARF10/16/17* ^68^, while miR167 antagonizes ARF6/8 ^47^. Our findings further illuminate the relatively unexplored role of miR160c in regulating *ARFs*, suggesting a mechanism by which it fine-tunes plant defense against phloem-feeding insects (Figure 6 and Figure 7).

Age-related changes in sRNA levels have been shown to play an important role in plant growth and stress responses. In Arabidopsis, the decline in the levels of miR156/7 with age drives plant maturation ^69,70^, while an increase in miR172 with plant age promotes the transition toward adulthood and flowering^71^. Notably, age-related miRNA gradients are also implicated in plant immunity against pathogens. For example, miR172b levels are very low during the early stage of seedling development but increase over time, enhancing pathogen immunity by reducing the expression of *TARGET OF EAT1/2* (*TOE1/2*), repressors of the immune receptor *FLAGELLIN-SENSING2* (*FLS2*) ^72^. In this study, we demonstrate the role of age-related sRNA changes in regulating insect resistance. Specifically, the decrease in miRNA160c levels with age alleviates inhibition of its target defense-related TFs, thereby promoting insect resistance (Figure 7).

The regulation of miRNA levels in plants is complex, and influenced by various factors including phytohormones, which can either increase or decrease miRNA levels ^19,73^. We found that auxin increased miR160c levels by promoting its transcription (Figure 7J and 7K). The connections between miRNAs and phytohormones are an emerging area of research, with multiple recent studies highlighting the role of miRNAs in regulating plant phytohormonal pathways ^73–75^. As a supplementary finding, we revealed that auxin-induced miRNA can target key regulators involved in the antagonism of SA synthesis (Figure S11). These findings broaden our understanding of the molecular and physiological mechanisms underlying phytohormone interactions. Our work demonstrates, for the first time, the role of miRNA in mediating the crosstalk between developmental auxin and defensive SA, balancing plant growth and defense (Figure S11).

Developing new crop varieties with improved product traits and enhanced stress resilience is a primary goal of global crop breeding, however, this effort is often limited by the growth-defense (G-D) trade-off ^75^. Consequently, the G-D trade-off has garnered increasing attention. However, the G-D balance in plants is not immutable; it can be influenced by various factors, such as developmental stage and environmental conditions. In this study, we specifically focused on the mechanisms underlying the trade-off between growth and insect resistance across different plant ages (Figure S11). We conclude that the auxin- miR160c module represses the ARFs-MYB-PAL module, mediating insect defense while ensuring early growth during the juvenile stage. As plants mature, this repression is gradually alleviated, allowing for the establishment of SA-mediated resistance alongside continued plant growth. These findings are likely to inspire further strategies for regulating crop growth and stress resistance by fine-tuning phytohormone interactions, key TFs, metabolic pathways, and miRNA levels. Importantly, genetic optimization in breeding programs must fully consider the age dynamics of plant-insect interactions to overcome the G-D trade-offs and develop ideal crop varieties.

## EXPERIMENTAL PROCEDURES

### Insect and plant materials

The *Bemisia tabaci* MEAM1 whitefly (*mtCOI* GenBank accession: GQ332577) was used in this study. Whiteflies were reared on cotton plants (*Gossypium hirsutum* cv. Zhemian 1793). Whitefly adults within 7 days post-emergence were used for experiments. Green peach aphids (*Myzus persicae*) were reared on tobacco (*N. tabacum* cv. NC89), with adult winged aphids also taken at 7 days post-emergence for experiments. The western flower thrips (*Frankliniella occidentalis*) were similarly reared on cotton plants (*G. hirsutum* cv. Zhemian 1793). Adult thrips, within 7 days post-emergence, were taken in the experiments.

*N. benthamiana* seeds (accession LAB and line H2B-RFP), tobacco (*N. tabacum* cv. NC89), and tomato (*Solanum lycopersicum* cv. Hezuo903) were sown in a soil mixture composed of peat moss, perlite, and vermiculite in a ratio of 5:3:1 (v/v/v) and placed in climate chambers. After approximately 14 days, when the *N. benthamiana*, tobacco, and tomato seedlings had germinated in a seed starter tray, the resulting plants were transplanted into plastic pots for individual cultivation. *N. benthamiana* plants at 7 days post-transplanting (21 days old, 3–4 true leaves) were classified as juvenile plants (low resistance), while *N. benthamiana* plants at 21 days post-transplanting (35 days old, 8–9 true leaves) were referred to as adult plants (high resistance) (Figure 2D). For tobacco, 7 days post-transplanting (21 days old, 3–4 true leaves) were classified as juvenile plants and 21 days post-transplanting (35 days old, 6–7 true leaves) were classified as adult plants. For tomatoes, 7 days post-transplanting (21 days old, 3–4 true leaves) were classified as juvenile plants and 21 days post-transplanting (35 days old, 8–9 true leaves) were classified as adult plants.

Insect-rearing plants and experimental plants were grown in climate chambers at Zhejiang University (Hangzhou, China) at 260 ±0 1°C, 65 0±0 10% relative humidity, and a photoperiod of 140 h light: 100 h dark. All the insect-plant interaction experiments were conducted under the same conditions.

### Insect bioassay

Bioassays were conducted using the following methods and standards. The test plants were placed in specially designed nylon cages (30 cm × 30 cm × 30 cm, 120 mesh), and the test insects (5 female and 5 male adults of MEAM1 whitefly, 3 female adults of aphid, and 5 female and 5 male adults of thrip respectively) were released into each of the cages. After two days, the performance of insects (whitefly, survival rate, and fecundity; aphid, offspring number; thrips, survival rate) was observed and recorded.

For the age-related bioassays, 21-day-old and 35-day-old *N. benthamiana* plants were simultaneously prepared based on their growth timeline. Specifically, we first sowed the initial batch of *N. benthamiana* and when these plants reached 14 days old, we sowed the second batch. Both batches were grown in a strictly controlled environment as described above. By the time the first batch reached 35 days, the second batch was 21 days old. These two batches of *N. benthamiana* were then used for subsequent bioassays as mentioned above. Bioassays related to gene and miRNA functions were performed through *Agrobacterium*-mediated transient gene expression or VIGS. 2 days after gene overexpressing or 14 days after gene silencing (Figure 2D), whiteflies were released onto the plants, and whitefly survival and fecundity were determined after two days. For bioassays involving the external application of chemicals to the plants, whitefly assessments commenced two days after chemical treatment.

### Transient expression assays

The full-length cDNAs of the selected genes (*NbPAL6*, *NbMYB42*, and *NbARF18Ls*) were amplified via PCR using gene-specific primer pairs (Table S2). The full-length CDS sequences were inserted into pCAMBIA1305 vectors using the NovoRec plus one-step PCR cloning kit (Novoprotein, Suzhou, China) to generate gene expression constructs (pCAMBIA1305-*NbPAL6*-GFP, pCAMBIA1305-*NbMYB42*-GFP, pCAMBIA1305-*NbARF18La*-GFP, and pCAMBIA1305-*NbARF18Lb*-GFP). For transient expression in *N. benthamiana*, *Agrobacterium tumefaciens* strain EHA105 containing the respective vector was cultured overnight in LB medium at 280°C. Bacterial cultures were harvested by centrifugation, resuspended in buffer (10 mM MES, pH 5.7, 10 mM MgCl_2_, and 200 µM acetosyringone) to an OD600 of 00.8, and infiltrated into leaves of 21-day-old or 35-day-old *N. benthamiana* using a needleless syringe (Figure 2D). The infiltrated plants were then maintained in the previously described climate chambers. Two days later, the treated plants were used for follow-up experiments including bioassays and sample collection (Figure 2D).

### VIGS assays

A 300 bp fragment of the target gene from *N. benthamiana* was identified using the VIGS tool on the Sol Genomics Network (https://vigs.solgenomics.net/). The gene fragment was amplified from *N. benthamiana* cDNA via PCR and cloned into the pTRV2 vector. The resulting plasmid was introduced into *A. tumefaciens* strain EHA105 by electroporation. Following cultivation, bacterial cells were harvested, resuspended, and adjusted to an OD600 of 0.2. The cultures were incubated at room temperature for at least 3 h. For leaf infiltration, cultures containing pTRV1 were mixed in equal volumes with those containing pTRV2:00 (empty vector, control), pTRV2-*NbPAL6*, pTRV2-*NbMYB42* or pTRV2-*NbARF18Ls*. These suspensions were infiltrated into 21-day-old *N. benthamiana* leaves using a 1 mL needleless syringe (Figure 2D). Following infiltration, the *N. benthamiana* plants were maintained in the same climate chamber conditions. Fourteen days later, the treated plants were used for follow-up experiments, including bioassays and sample collection (Figure 2D).

### Subcellular localization

The pCAMBIA1305-*NbMYB42*-GFP, pCAMBIA1305-*NbARF18La*-GFP, pCAMBIA1305-*NbARF18Lb*-GFP, and pCAMBIA1305-GFP (empty vector) constructs were used for subcellular localization. These constructs were infiltrated into the leaves of *N. benthamiana* line H2B-RFP via *A. tumefaciens* strain EHA105. Two days post-infiltration, GFP fluorescence signals were visualized using confocal microscopy (Zeiss LSM710, Oberkochen, Germany).

### RNA extraction and RT-qPCR analysis

Total RNA was extracted using AG RNAex Pro reagent (Accurate Biology, Changsha, China). For mRNA, single-stranded cDNA was synthesized using the *Evo M-MLV* RT Kit with gDNA Clean for qPCR Ver.2 (Accurate Biology Changsha, China). For plant miRNA cDNA synthesis, the miRNA 1st strand cDNA synthesis kit (Accurate Biology, Changsha, China) was used. RT-qPCR was conducted on a Bio-Rad CFX96 real-time PCR system (Bio-Rad, California, USA) with specific primers (Table S2). Data were analyzed with Bio-Rad CFX Manager Software, employing the 2^-ΔΔC^ method. The *N. benthamiana* housekeeping gene *NbACTIN* served as the internal control for gene expression analysis, and the expression of the *NbU6* snRNA genes was employed as the internal control for miRNA level analysis.

### Measurement of phytohormone content

For the quantification of phytohormones (SA, JA, and IAA), *N. benthamiana* leaves were promptly excised and ground into powder in liquid nitrogen. Leaf powder (0.15 g) was mixed with 1 mL of HPLC-grade ethyl acetate (Sinopharm, Shanghai, China) containing 10 ng D4-SA (Quality Control Chemicals, Spokane, WA, USA), 10 ng D6-JA (Quality Control Chemicals, Spokane, WA, USA), and 5 ng D5-IAA (OlhemIm, Shanghai, China). The mixture was fully vortexed and centrifuged. The supernatant was collected and dried using a vacuum concentrator (Eppendorf, Hamburg, Germany) at 30°C. The residue was resuspended in 110 µL of methanol:H_2_O (50:50, v/v), and the resulting supernatant was analyzed via high-performance liquid chromatography-tandem mass spectrometry (TripleTOF 5600+; AB Sciex, Redwood City, CA, USA).

### Degradome sequencing

The degradome sequencing analysis, performed according to established protocols from a previous study ^76^, was outsourced to Lianchuan Biotechnology Co., Ltd. Poly(A) RNA was purified from total plant RNA using poly-T oligo magnetic beads (Invitrogen, Carlsbad, CA, USA) through two rounds of purification.

First-strand cDNA synthesis was performed with a 3’-adapter random primer, followed by size selection using AMPureXP beads (Beckman Coulter, Indianapolis, IN, USA) and PCR amplification. The final cDNA library had an average insert size of 200–400 bp. Sequencing was carried out with 50 bp single-end reads on an Illumina HiSeq 2500 (Illumina, San Diego, CA, USA). CleaveLand 3.0 and TargetFinder 50 were employed to identify potential sliced targets of known miRNAs.

### Y1H and Y2H assay

For the Y1H assay, approximately 2 0kb of the promoter regions of the target genes (*NbPAL6* and *NbMYB42*) were inserted into the pAbAi vector to construct the bait plasmids. These plasmids were integrated into the Y1HGold yeast strain via polyethylene glycol-mediated transformation, following in the Yeast User Manual PT4087-1 (Clontech, Shiga, Japan). The CDSs of the corresponding TFs *NbMYB42* and *NbARF18La/b* were cloned into the pGADT7 vector. The relevant vectors were co-expressed in Y1H yeast cells and screened with varying concentrations of Aureobasidin A (AbA, TaKaRa, Beijing, China).

For the Y2H assays, the full-length CDSs of the target genes *NbARF18La* and *NbARF18Lb* were ligated into pGBKT7 and pGADT7 (Clontech, Shiga, Japan), respectively. The paired vectors were transformed into Y2HGold yeast strain cells via PEG/LiAc transformation and grown on SD −Leu/−Trp medium. After 3–4 days of incubation, the clones were transferred onto SD −Leu/−Trp/−His/−Ade medium and incubated at 29°C for 3–4 days. The primers used for vector construction related to the Y1H and Y2H assays are listed in Table S2.

### Histochemical GUS activity assays

For the reporter constructs, the promoters of the target genes *NbPAL6* and *NbMYB42* were introduced into the pBI121 vector containing a *GUS* gene-coding region. Simultaneously, the full-length CDS of the corresponding target TFs (*NbMYB42* and *NbARF18La/b*) were inserted into the pCAMBIA1305 vector to generate effector constructs (pCAMBIA1305-*NbMYB42*-GFP, pCAMBIA1305-*NbARF18La*-GFP, pCAMBIA1305-*NbARF18Lb*-GFP, as described in the Transient expression assays section). The pCAMBIA1305 vector without the target CDS was used as the negative control (empty effector). These vectors were transformed into *A. tumefaciens* strain EHA105, and reporter vectors were co-transformed with the corresponding effector or control vectors for transient expression. GUS activity was assessed using a GUS histochemical assay kit (Coolaber, Beijing, China) according to the manufacturer’s instructions. GUS expression was visualized through staining and documented using an iPhone 13 Pro Max (Apple, California, USA).

### Dual-luciferase assays

For the reporter construct, the promoters of the target genes (*NbPAL6* and *NbMYB42*) were inserted into the reporter vector pGreenII 0800-LUC. Effector constructs, including pCAMBIA1305-*NbMYB42*-GFP, pCAMBIA1305-*NbARF18La*-GFP, pCAMBIA1305-*NbARF18Lb*-GFP, and the empty vector pCAMBIA1305, were the same as those used in the Histochemical GUS activity assays. These vectors were transformed into *A. tumefaciens* EHA105 (pSoup). In this system, *LUC* expression was driven by the target gene promoters, while *REN* was driven by the CaMV *35S* promoter as an internal control. The reporter and corresponding effector constructs were transiently transformed into *N. benthamiana* leaves. After a three-day incubation, LUC signals were measured using the Dual-Luciferase Reporter Assay Kit (Vazyme, Nanjing, China) and analyzed on a FlexStation 3 Multi-Mode Microplate Reader (Molecular Devices, San Jose, CA, USA). Relative luciferase activity was calculated based on LUC:REN ratios.

### ChIP assay

Leaves transformed with pCAMBIA1305-*NbMYB42*-GFP, pCAMBIA1305-*NbARF18La*-GFP, or pCAMBIA1305-*NbARF18Lb*-GFP (as described in the Transient expression assay), were collected for ChIP assays. The experiment followed established protocols using the same transient gene expression system ^77^. Briefly, transformed *N. benthamiana* leaves were harvested and fixed with 1% formaldehyde. The fixation was quenched by adding 1 × PBS buffer containing 0.125 M glycine under vacuum for 5 min. Chromatin DNA was sonicated to fragments of approximately 200–300 bp. The chromatin complexes were immunoprecipitated (IP) by Protein G Agarose (Beyotime, Shanghai, China) with an anti-GFP monoclonal antibody (Beyotime, Shanghai, China). The co-IP DNA was recovered at 65°C, purified, and analyzed by qPCR with SYBR qPCR Mix (Accurate Biology, Changsha, China) with gene-specific primers (Table S2). The relative fold enrichment was calculated using leaves transformed with the pCAMBIA1305-GFP (empty vector) as a control.

### EMSA assay

To prepare the test TF proteins for EMSA, the CDS of *NbMYB42* was fused into the GST-tagged pGEX- 6P-1 vector to generate NbMYB42-GST. The *NbARF18La* CDS was fused into the His-tagged pET28a- SUMO vector, generating NbARF18La-His-SUMO, and the *NbARF18Lb* CDS was fused into the MBP-tagged pMAL-c5x vector, generating NbARF18Lb-MBP. These fusion proteins were expressed in *E. coli* BL21 cells using 0.2 mM isopropyl-β-D-thiogalactopyranoside (IPTG) and purified. For protein purification, NbMYB42-GST was purified through GST-affinity chromatography (Beyotime, Shanghai, China), NbARF18Lb-MBP using MBPSep Dextrin Agarose Resin 6FF (Yeasen, Shanghai, China) and NbARF18La-His with Ni NTA Beads 6FF (Smart-Lifesciences, Changzhou, China).

Putative binding sites in the target gene promoters were predicted by Jasper (https://jaspar.elixir.no/). For the *NbMYB42* and *NbPAL6* promoter interaction, biotin-labeled probes were designed based on the P3 sequence (GGTGGTTGTTGAGAGG) of the *NbPAL6* promoter (Figure 3G; Table S2), with a mutated version (AAAAAAAAAAAAAAAA) serving as the mutant probe. For the interaction between NbARF18La/b and the *NbMYB42* promoter, biotin-labeled probes targeting the AuxRE element P2 sequence (TGTCTC) were designed (Figure 4G and 4H; Table S2), with a mutated version (AAAAAA) as the control. Cold competitors (10-, 20-, and 50-fold excess) of unlabeled probes were included in the reactions. Probes were labeled using the EMSA Probe Biotin Labeling Kit (Beyotime, Shanghai, China). Approximately 300 ng of the fusion protein was incubated with 30 ng of labeled probes, and specific binding was assessed via the Chemiluminescent EMSA Kit (Beyotime, Shanghai, China) following the manufacturer’s instructions. Chemiluminescent imaging was performed with the ChemiDoc Touch system (Bio-Rad, California, USA). Labeled probes incubated with GST, His-SUMO, and MBP proteins served as negative controls.

### Pull-down assay

Pull-down assays were conducted to determine the interaction of NbARF18La and NbARF18Lb. The recombinant proteins NbARF18La-His and NbARF18Lb-MBP obtained in the above-mentioned EMSA assay were further used in this test. For MBP pull-down, MBPSep Dextrin Agarose Resin (Yeasen, Shanghai, China) is prepared by washing them with PBS and then incubating them with MBP-bait protein NbARF18Lb or control MBP protein. After washing, the beads are incubated with His-tagged interacting protein NbARF18La at 4°C for 3 h. Then the MBP beads underwent four washes with washing buffer (20 mM phosphate buffer, 500 mM NaCl, 100 mM imidazole, pH 7.4) and were then collected using collecting buffer (20 mM phosphate buffer, 500 mM NaCl, 500 mM imidazole, pH 7.4). The eluted samples were separated by SDS-PAGE and detected by an anti-MBP (Affinity Bioscience, Jiangsu, China) antibody and an anti-His antibody (Proteintech, Wuhan, China).

### Bimolecular fluorescence complementation (BiFC) assay

The CDSs of *NbARF18La* and *NbARF18Lb* were inserted into p2YN and p2YC vectors, respectively. *A. tumefaciens* EHA105 cells carrying p2YN constructs were combined with cells containing p2YC constructs at a 1:1 ratio. This mixture was then infiltrated into the leaves of the *N. benthamiana* H2B-RFP line. After three days of incubation, fluorescence was visualized under a laser confocal microscope (Zeiss LSM710, Oberkochen, Germany).

### Exogenous application of phytohormones

The target compound was dissolved in methanol (for SA, IAA, and NAA) or dimethyl sulfoxide (DMSO) (for Yucasin and _L_-Kyn) to produce the stock solution. Lanolin was selected as a slow-release medium. The lanolin was heated to a liquid state and the stock solution was added to the lanolin to create the final mixture: SA at 1 mM; IAA and NAA at 1, 25, and 50 μM; Yucasin and _L_-Kyn at 50 μM. All the compounds were purchased from Aladdin, Shanghai, China. The lanolin solution containing the target compounds was transferred to a 1 mL sterile syringe and stored at -20°C.

Exogenous SA was applied to 21-day-old and 35-day-old *N. benthamiana* plants. IAA and NAA were applied to 35-day-old plants, while Yucasin and _L_-Kyn were applied to 21-day-old plants. For application, 0.25 mL of lanolin was squeezed from the syringe and gently smeared along the midvein at the base of the leaves using latex-gloved fingers. After two days (Figure 2D), the treated plants were used in bioassays or for sample collection for further experiments.

### Transient expression and functional inhibition of miRNA in N. benthamiana

To overexpress target miRNA *Nb*mi160c in *N. benthamiana*, the precursor sequences of *Nb*miR160c were synthesized by GenScript (Nanjing, China) and inserted into the pCAMBIA1300 vector through specific primers (Table S2; Figure S8A). The constructs were infiltrated into 35-day-old *N. benthamiana* plants, and after two days (Figure 2D), the treated plants were used in bioassays or for sample collection for further experiments.

To inhibit the *Nb*miR160c function, a target mimicry (mimic160c) expression system was introduced.

The precursor sequences of mimic160c were synthesized by GenScript (Nanjing, China) and inserted into the pCAMBIA1300 vector using specific primers (Table S2; Figure S8F). The constructs were infiltrated into 21-day-old *N. benthamiana* plants, and after two days (Figure 2D), the treated plants were used for bioassays or sample collection for further experiments.

### MiRNA targeting validation

GFP, GUS, and LUC reporter systems were used to investigate the cleavage of *NbARF18La* and *NbARF18Lb* targets by *Nb*miR160c. For the GFP reporter, the *Nb*miR160c target site of *NbARF18La* and *NbARF18Lb* mRNA, as well as its mutant form (Figure S8C) were cloned into the pCAMBIA1305-GFP vector to construct GFP sensors through specific primers (Table S2; Figure S8D). The constructs of the target site and mutant target site were co-infiltrated with pCAMBIA1300-*Nb*miR160c through *A. tumefaciens* strain EHA105. After two days of infiltration, GFP fluorescence was observed using confocal microscopy, as described above. For the GUS and LUC reporters, the *Nb*miR160c target site of *NbARF18La* and *NbARF18Lb* mRNA, along with its mutant form, were cloned into the pCAMBIA121-GUS and pGreenII 0800-LUC vectors, respectively, to construct GUS and LUC sensor through specific primers (Table S2; Figure S8D). These constructs were co-infiltrated with pCAMBIA1300-*Nb*miR160c. After two days of infiltration, GUS and LUC activity was examined using the previously described method.

### MiRNA dual-luciferase assay

Co-expression of the LUC reporter and *Nb*miR160c was carried out to evaluate the influence of *Nb*miR160c on downstream genes. The effectors pCAMBIA1300-*pre*-*Nb*miR160c, pCAMBIA1300- premimic160c, and pCAMBIA1300 (empty vector control) were co-expressed with LUC reporters linked to the *NbMYB42* or *NbPAL6* promoters. After two days of infiltration, LUC activity was examined using the method mentioned method. The combination of effectors and reporters is shown in Figure S9A to S9D.

The system was also used to evaluate the possible influence of *Nb*miR160c on *NbPAL6* transcription when *NbARFs* were overexpressed. The *LUC* gene linked to the *NbPAL6* promoter or *NbMYB42* promoter was used as the reporter respectively. pCAMBIA1305-*NbARFs* plus pCAMBIA1300-*pre*- *Nb*miR160c, and pCAMBIA1305-*NbARFs* plus pCAMBIA1300 (empty vector) were co-infiltrated with the reporter into the leaves of *N. benthamiana*. LUC activity for the different combinations was detected and compared. The combinations of effectors and reporters are shown in Figure 7G and Figure S9E.

### Statistical analysis

All data were analyzed using IBM SPSS Statistics 22.0 and visualized using GraphPad Prism 9. Statistical significance was determined using Student’s *t*-test (two-tailed) or one-way ANOVA, followed by Fisher’s least significant difference (LSD) test.

## RESOURCE AVAILABILITY

### Lead contact

Further information and requests for resources and reagents should be directed to and will be fulfilled by the lead contact, Xiao-Wei Wang.

### Materials availability

This study did not generate any new unique reagents or materials to report. All reagents or materials used are commercially available.

### Data and code availability

Any additional information required to reanalyze the data reported in this paper is available from the lead contact upon request.

## ACKNOWLEDGMENTS

We thank Rong Jin from the Agricultural Experiment Station, Zhejiang University, for greenhouse management. We also thank Xiao-Dan Wu from the Analysis Center of Agrobiology and Environmental Sciences, Zhejiang University, for technical support in the detection and analysis of phytohormones. X.-W.W. was supported by the National Natural Science Foundation of China (31925033, 32161143008) and the National Key Research and Development Program (2022YFC2601002).

## AUTHOR CONTRIBUTIONS

Conceptualization, W.-H.H., F.-B.Z., J.-X.W. and X.-W.W.; methodology, F.-B.Z., W.-H.H., and X.-P.F.; Investigation, F.-B.Z., W.-H.H., S.-X.J., J.-X.W., X.-P.F., and K.-L.L.; writing—original draft, W.-H.H. and F.-B.Z.; writing—review & editing, W.-H.H., F.-B.Z., S.-X.J. and X.-W.W.; funding acquisition, X.-W.W.; supervision, S.-S.L., and X.-W.W..

## SUPPLEMENTAL INFORMATION

**Document S1. Figures S1–S11 and Table S1-S2**

