## Supplementary Figures and Tables for "Auxin-salicylic acid seesaw regulates the age-dependent balance between plant growth and herbivore defense"

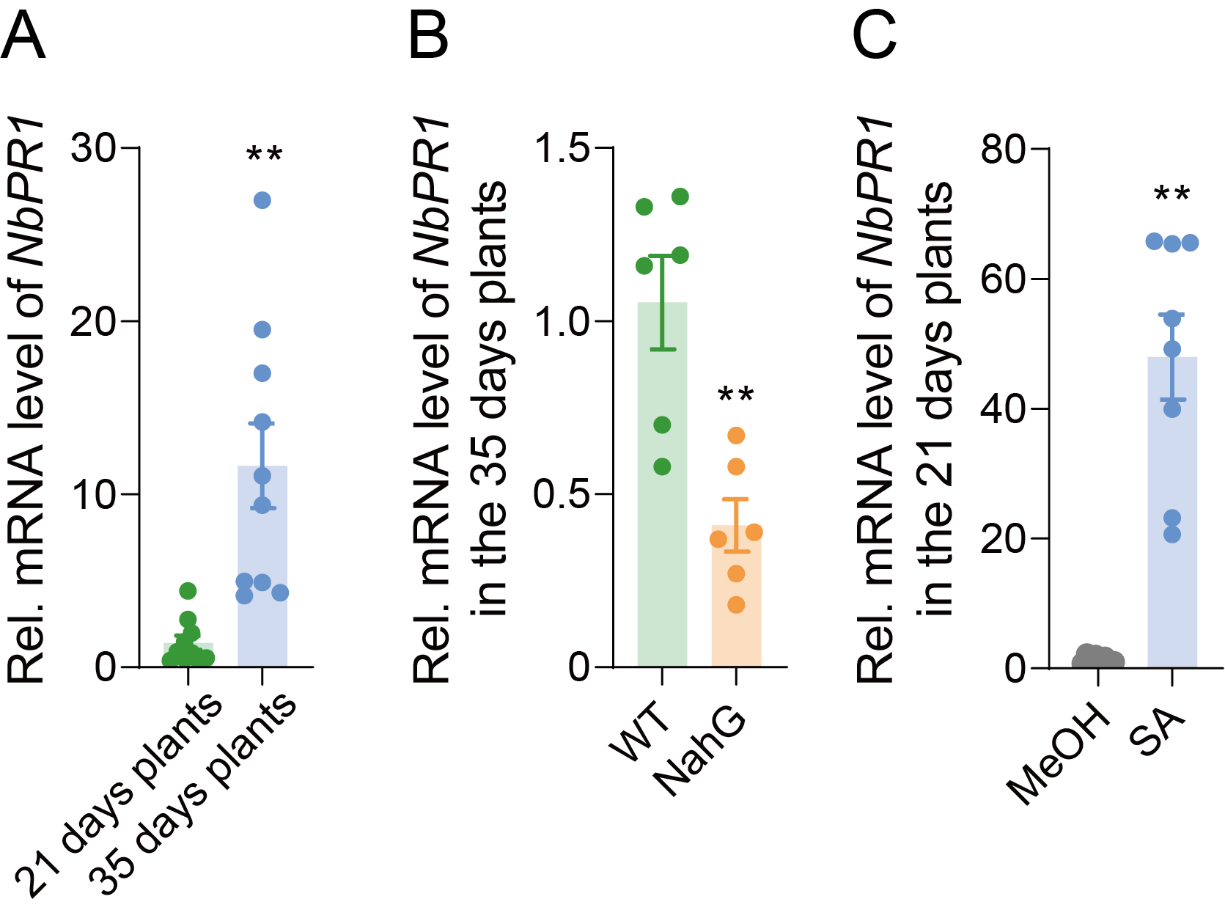

**Figure S1. Expression of the *NbPR1* gene downstream of SA at different ages and SA levels**

(A) Expression level of the SA downstream gene *NbPR1* in the *N. benthamiana* plants at 21 and 35 days old. (B) Expression level of *NbPR1* in 35-day-old wild-type and NahG *N. benthamiana* plant. (C) Expression level of *NbPR1* in 21-day-old *N. benthamiana* plants treated with 1 mM SA. Values are mean ± SE, n = 10 for A n = 6 for B; n = 8 for C. Student’s *t*-test (two-tailed) was used for significant difference analysis. ***P* < 0.01.

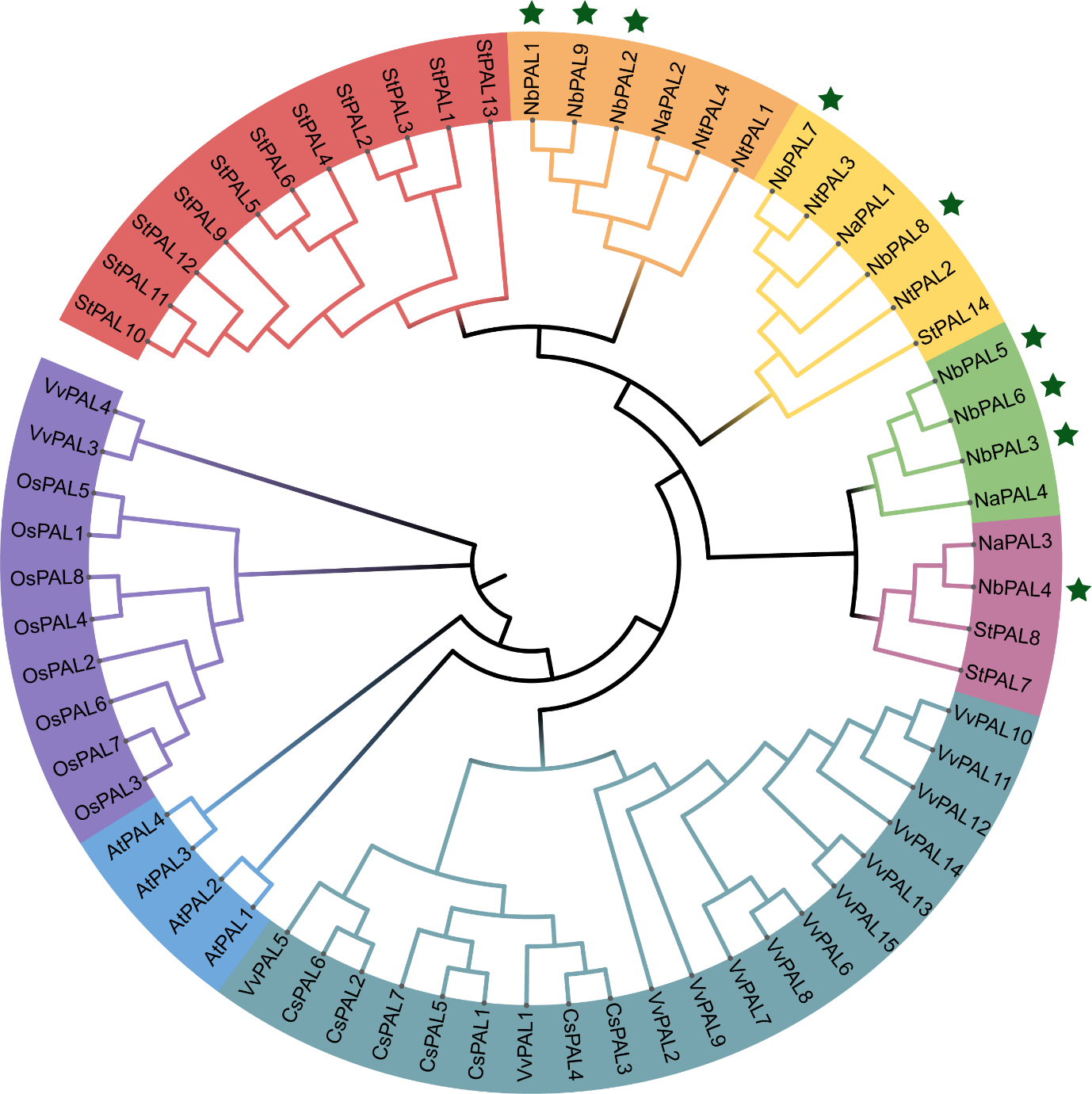

**Figure S2. Phylogenetic tree of PAL protein**

Amino acid sequences of PAL proteins from *N. benthamiana* (Nb), *N. tabacum* (Nt), *N. attenuata* (Na), *Arabidopsis thaliana* (At), *Oryza sativa* (Os), *Solanum tuberosum* (St), *Vitis vinifera* (Vv), and *Camellia sinensis* (Cs) were aligned by ClustalW. The Maximum Likelihood tree was constructed with MEGA X with 1000 bootstrap replicates.

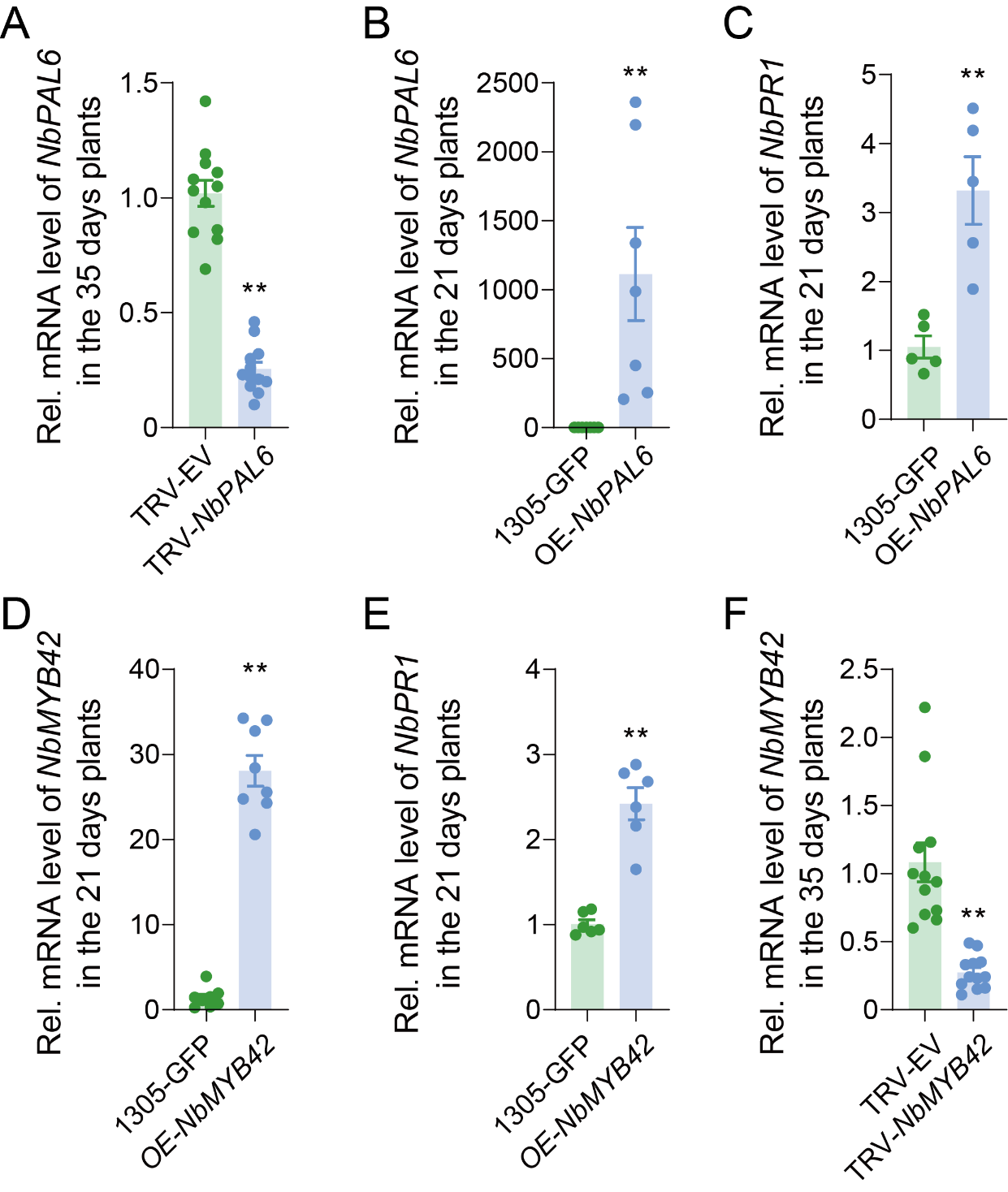

**Figure S3. Gene expression level after silencing and overexpressing *NbPAL6* and *NbMYB42***

(A) Expression level of *NbPAL6* in *NbPAL6*-silenced 35-day-old *N. benthamiana* plants. (B) Expression level of *NbPAL6* in *NbPAL6*-overexpressed 21-day-old *N. benthamiana* plants. (C) Expression level of SA downstream *NbPR1* gene in *NbPAL6*-overexpressed 21-day-old *N. benthamiana* plants. (D) Expression level of *NbMYB42* in the *NbMYB42*-overexpressed 21-day-old *N. benthamiana* plants. (E) Expression level of SA downstream *NbPR1* gene in the *NbMYB42*-overexpressed 21-day-old *N. benthamiana* plants. (F) Expression level of *NbMYB42* in *NbMYB42*-silenced 35-day-old *N. benthamiana* plants. Values are mean ± SE, n = 12 for A and F; n = 7 for B; n = 5 for C; n = 8 for D; n = 6 for E. Student’s *t*-test (two-tailed) was used for significant difference analysis. ***P* < 0.01.

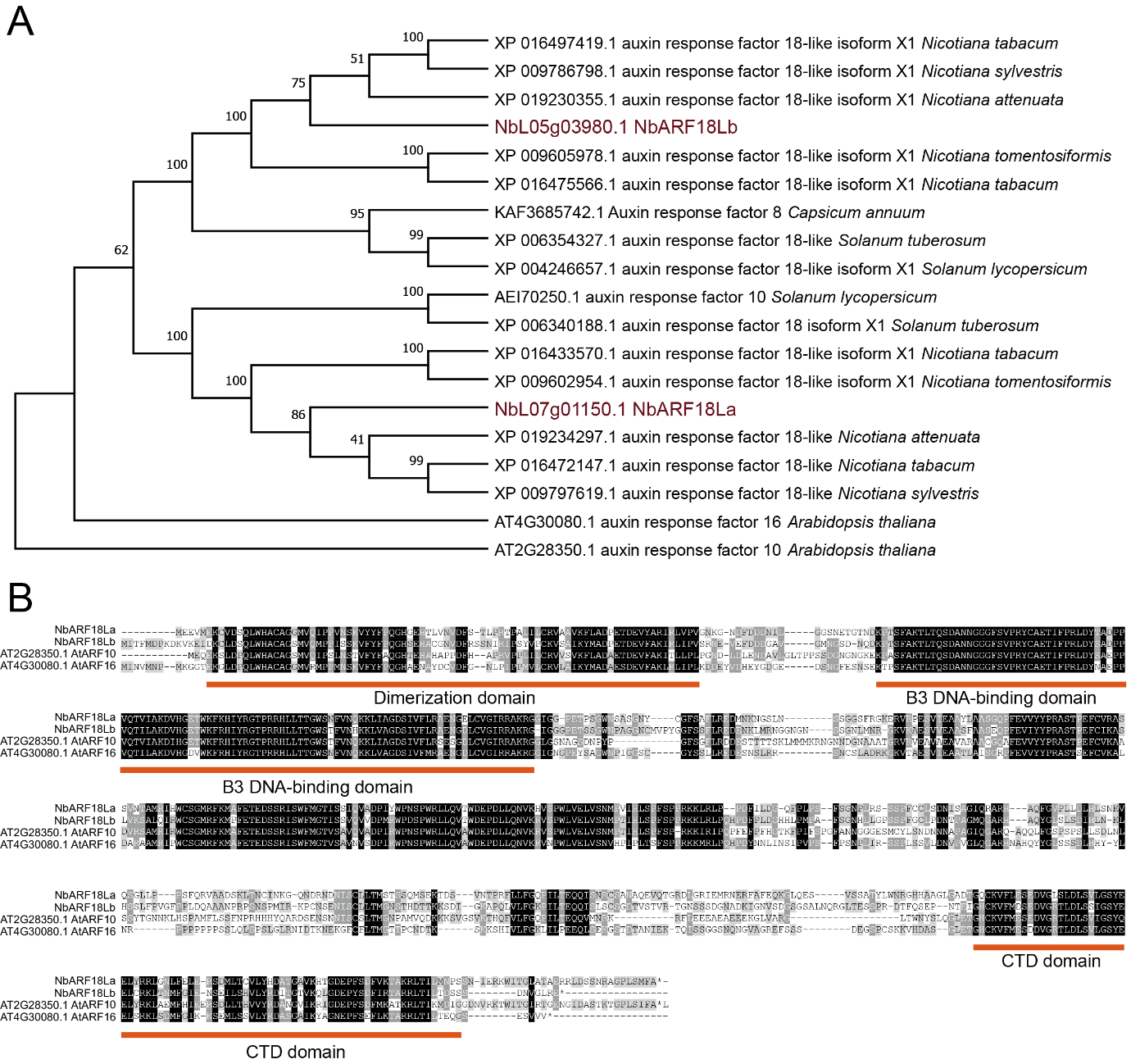

**Figure S4. Phylogenetic relationship and conserved DNA binding domain of NbARF18La/b**

(A) Phylogenetic tree of ARF proteins. Homologous genes of *NbARF18Ls* were identified using NCBI Blast (https://blast.ncbi.nlm.nih.gov/Blast.cgi). Protein sequences were aligned using ClustalW, and a phylogenetic tree with 1000 bootstrap replicates was constructed by the Maximum Likelihood method in MEGA X. (B) Multi-sequence alignment analysis of ARF proteins. The B3 DNA binding domain (DBD) and C-terminal dimerization domain (CTD) regions are underlined. Protein sequence alignment results were visualized by GenDoc.

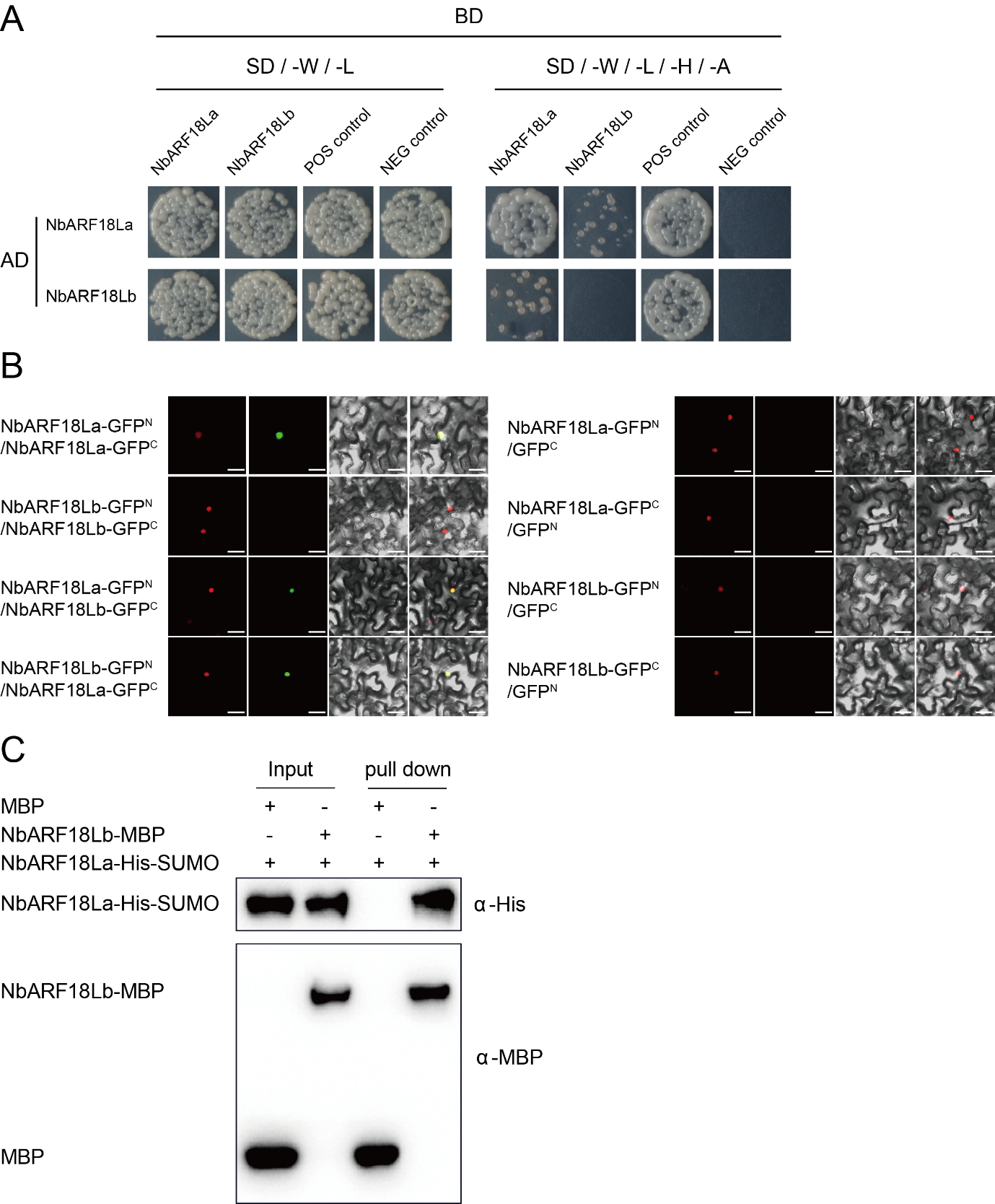

**Figure S5. NbARF18La interacts with NbARF18Lb**

(A) Yeast two-hybrid (Y2H) assay between NbARF18La and NbARF18Lb. Yeast cells co-transformed with NbARF18La and NbARF18Lb plasmids grow on a selective medium (SD/-W-L-H-A) for 3 days, indicating an interaction between NbARF18La and NbARF18Lb. Additionally, yeast cells co-transformed with NbARF18La plasmids (pGADT7-NbARF18La and pGBKT7-NbARF18La) also grew on selective medium (SD/-W-L-H-A), showing that NbARF18La exhibits self-interaction. (B) Bimolecular fluorescence complementation (BiFC) assays confirmed the interaction between NbARF18La and NbARF18Lb in *N. benthamiana* epidermal cells. BiFC assays show that NbARF18La interacts with NbARF18Lb in nuclei, and NbARF18La interacts with itself in nuclei. The BiFC signal (green fluorescence) observed in the nuclei indicates an interaction between NbARF18La and NbARF18Lb, as well as a self-interaction of NbARF18La. Confocal microscopy was used to visualize the fluorescence. Scale bars, 40 μm. (C) The interaction between NbARF18La and NbARF18Lb was further validated by an in vitro pull-down assay. Recombinant NbARF18Lb-MBP and NbARF18La-His-SUMO proteins were incubated with MBP beads in a pull-down binding buffer. The pulled-down proteins were detected by immunoblotting using anti-MBP and anti-His antibodies. A mixture of MBP and NbARF18La-His-SUMO was used as a negative control.

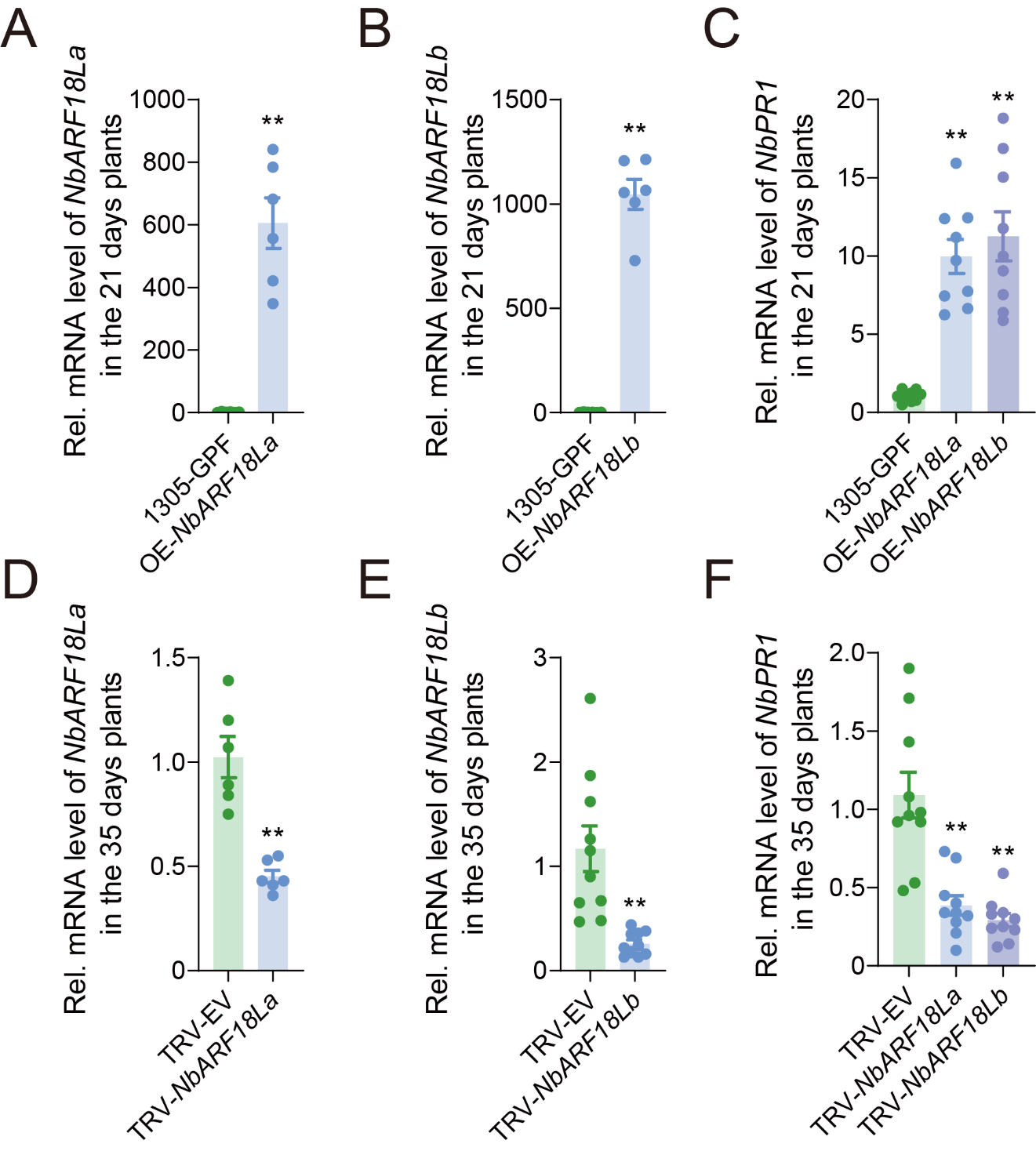

**Figure S6. Gene expression levels after overexpressing and silencing *NbARF18La/b***

(A, B) Expression levels of *NbARF18La/b* in the *NbARF18La/b*-overexpressed 21-day-old *N. benthamiana* plants. (C) Expression level of SA downstream gene *NbPR1* in the *NbARF18La/b*-overexpressed 21-day-old *N. benthamiana* plants. (D, E) Expression levels of *NbARF18La* and *NbARF18Lb* in the *NbARF18La/b*-silenced 35-day-old *N. benthamiana* plants. (F) Expression level of SA downstream gene *NbPR1* in the *NbARF18La/b*-silenced 35-day-old *N. benthamiana* plants. Values are mean ± SE, n = 6 for A, B and D; n = 9 for C; n = 10 for E and F. Student’s *t*-test (two-tailed) was used for significant difference analysis. ***P* < 0.01.

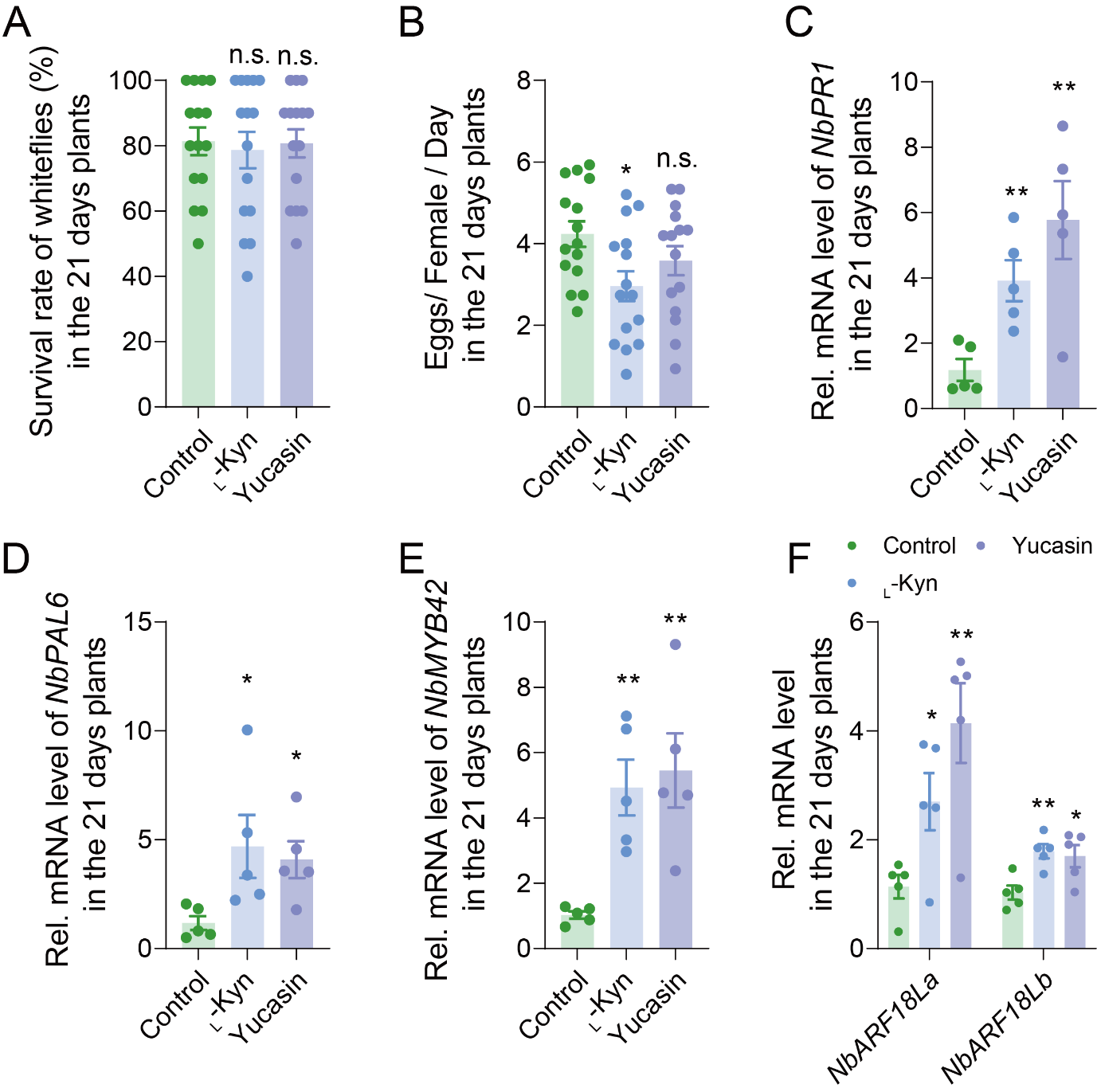

**Figure S7. Auxin synthesis inhibitors enhance the NbARF18La/b-MYB42 module in juvenile plants**

Survival rate and fecundity of whiteflies on N. benthamiana plants treated with _L_-Kyn and Yucasin (50 μM) at 21 days old. (B) SA levels in *N. benthamiana* plants treated with the IAA inhibitors _L_-Kyn and Yucasin (50 μM) at 21 days old. (C) Expression level of SA downstream gene *NbPR1* in _L_-Kyn- and Yucasin-treated (50 μM) 21-day-old *N. benthamiana* plants. (D) Expression level of *NbPAL6* in _L_-Kyn- and Yucasin-treated (50 μM) 21-day-old *N. benthamiana* plants. (E) Expression level of *NbMYB42* in _L_-Kyn- and Yucasin-treated (50 μM) 21-day-old *N. benthamiana* plants. (F) Expression levels of *NbARF18La/b* in the _L_-Kyn- and Yucasin-treated (50 μM) 21-day-old *N. benthamiana* plants. Values are mean ± SE, n = 15 for A and B; n = 5 for C, D, E, and F. Student’s *t*-test (two-tailed) was used for significant difference analysis. n.s., not significant; **P* < 0.05, ***P* < 0.01.

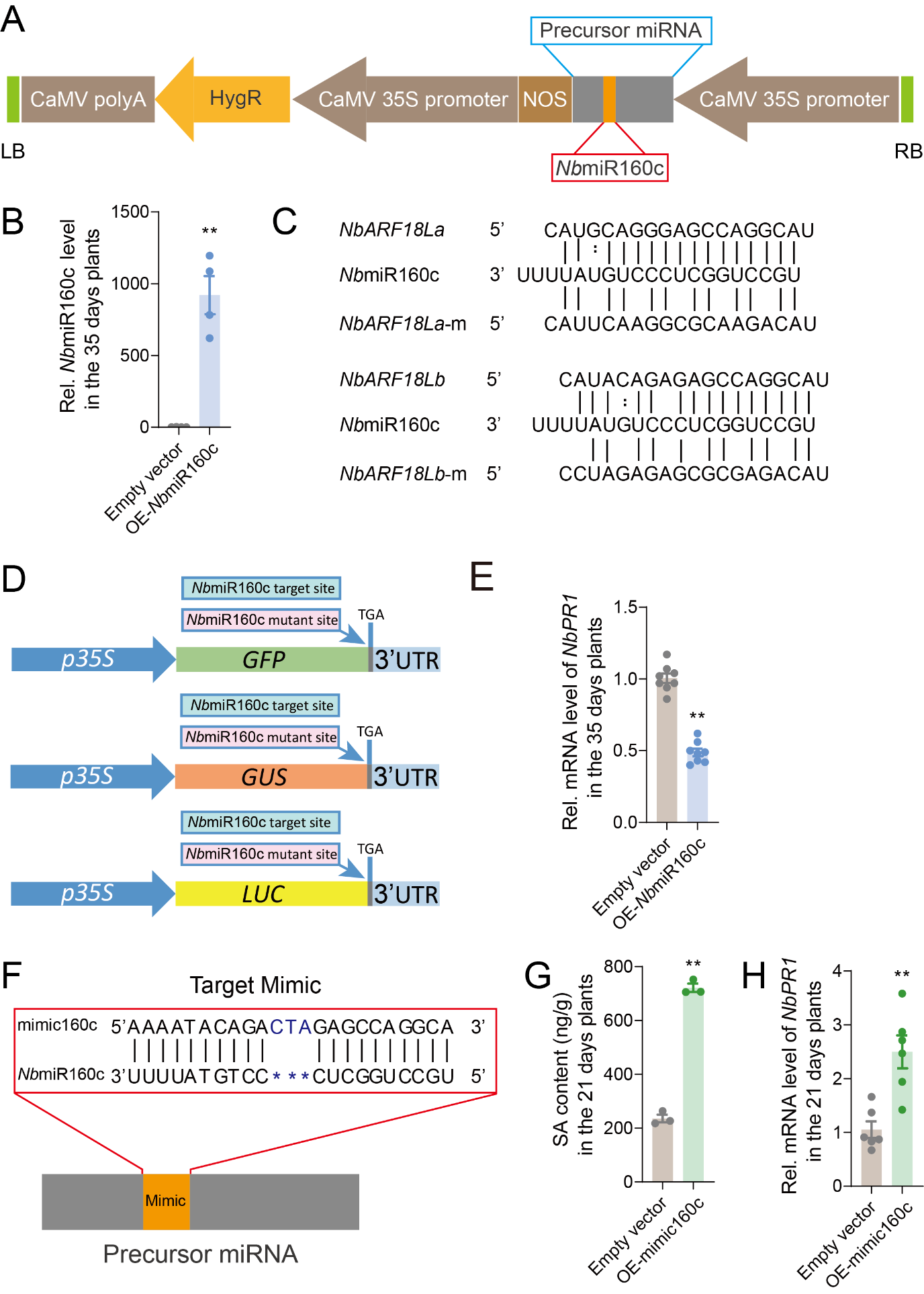

**Figure S8. Overexpression of mimicRNA160c increased the SA levels in juvenile plants**

(A) Schematic diagram of the *Nb*miR160c expression cassette *35S*-*Nb*miR160c. This vector was used to overexpress *Nb*miR160c in *N. benthamiana* plants through *Agrobacterium*-mediated transformation. The blue box represents the miRNA precursor, and the red box represents *Nb*miR160c. HygR: Hygromycin B phosphotransferase; NOS: Nos Terminator; RB: Right border; LB: Left border. (B) Expression level of *Nb*miR160c in the *Nb*miR160c-expressed 35-day-old plants. (C) Sequence alignment of *Nb*miR160c with the predicted target sites in *NbARF18La* and *NbARF18Lb*, along with their mutated versions (*NbARF18La*-m and *NbARF18Lb*-m) used in subsequent experiments. Colons represent G:U pairs. (D) Schematic diagram of the GFP/GUS/LUC sensor carrying the *Nb*miR160c target site and the mutant target site. TGA: Termination codon; UTR: Untranslated regions. (E) Expression level of *NbPR1* in the *Nb*miR160c-expressed 35-day-old plants. (F) Design of mimicry sequences (mimic160c) for *Nb*miR160c. (G) SA levels in mimic160c-expressed 21-day-old plants. (H) Expression level of *NbPR1* in the mimic160c-expressed 21-day-old plants. Values are mean ± SE, n = 4 for B; n = 8 for E; n = 3 for G; n = 6 for H. Student’s *t*-test (two-tailed) was used for significant difference analysis. ***P* < 0.01.

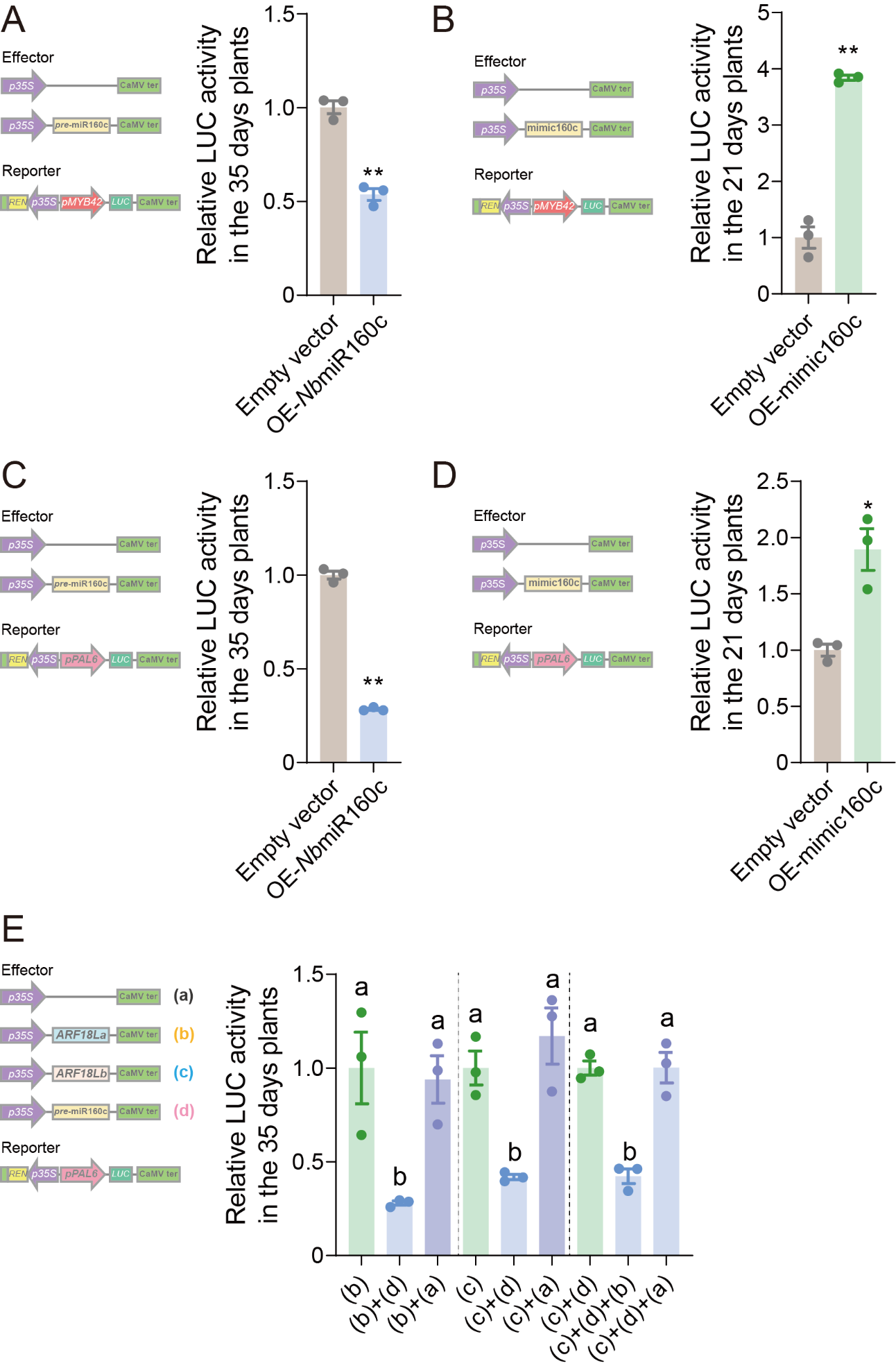

**Figure S9. *Nb*miR160c affects the transcription of *NbMYB42* and *NbPAL6***

(A to E) Schematic diagram showing the effector and reporter vectors used in transient transcriptional activity assays in *N. benthamiana* leaves. The data show the relative LUC activity normalized to REN activity. Values are mean ± SE, n = 3. Student’s *t*-test (two-tailed) was used for significant difference analysis in A, B, C, and D. **P* < 0.05, ***P* < 0.01. One-way ANOVA followed by Fisher’s least significant difference (LSD) test was used for significant difference analysis in E. Bars with different lowercase letters indicate significant differences between treatments at *P* < 0.05.

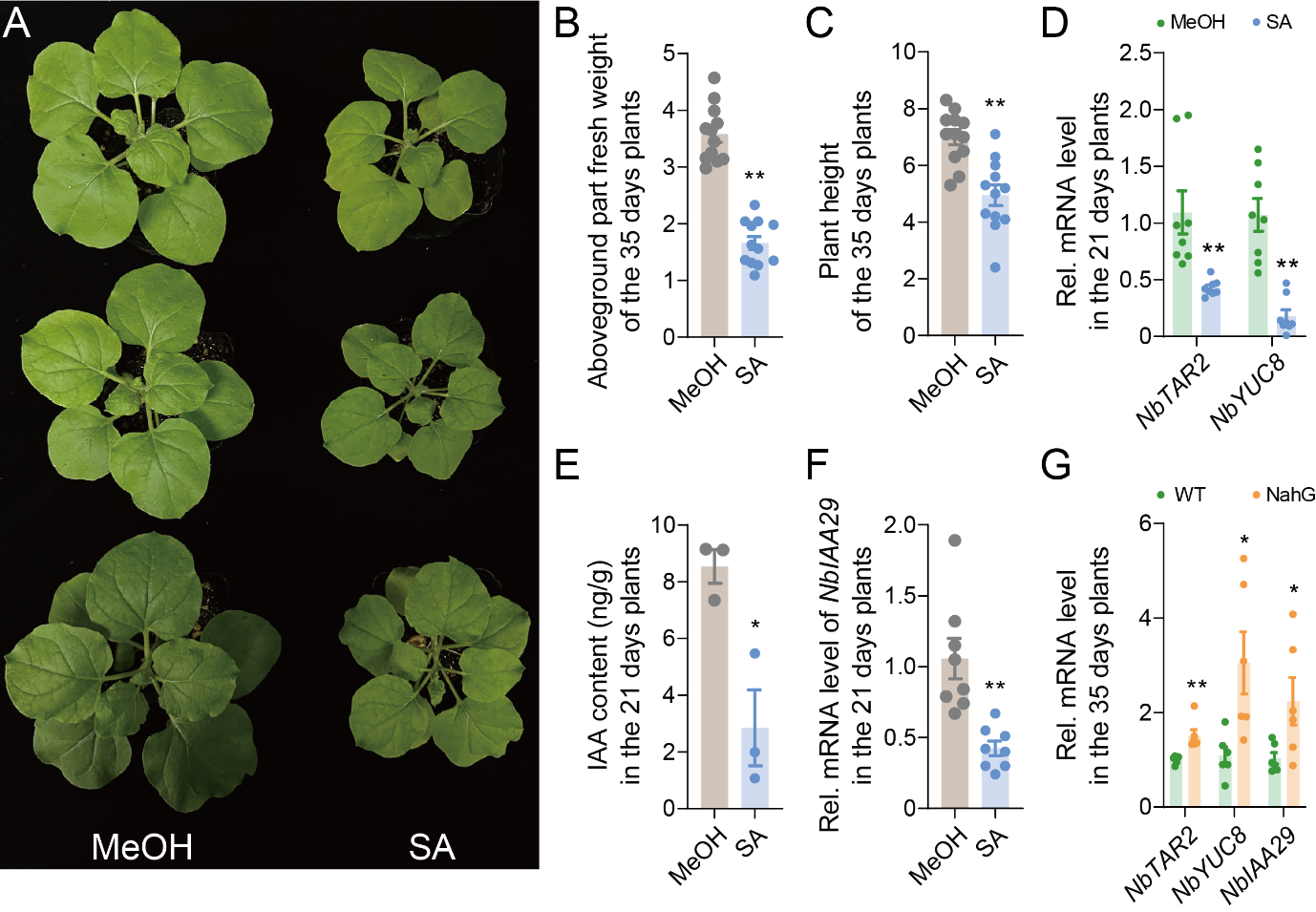

**Figure S10. Excessive SA impacts early plant growth by antagonizing auxin**

(A) Photographs of 35-day-old *N. benthamiana* plants treated with SA starting at 21 days old. (B, C) Aboveground fresh weight (B) and plant height (C) of SA-treated (1 mM) *N. benthamiana* plants. (D) Expression level of auxin synthesis genes *NbTAR2* and *NbYUC8* in 21-day-old *N. benthamiana* plants treated with 1 mM SA. (E) Auxin level in 21-day-old *N. benthamiana* plants after successive SA treatment (1 mM). (F) Expression level of auxin downstream gene *NbIAA29* in SA-treated (1 mM) 21-day-old *N. benthamiana* plants. (G) Expression level of auxin pathway genes in 35-day-old wild type and NahG *N. benthamiana* plants. Values are mean ± SE, n = 12 for B and C; n = 8 for D and F; n = 3 for E; n = 6 for G. Student’s *t*-test (two-tailed) was used for significant difference analysis. **P* < 0.05, ***P* < 0.01.

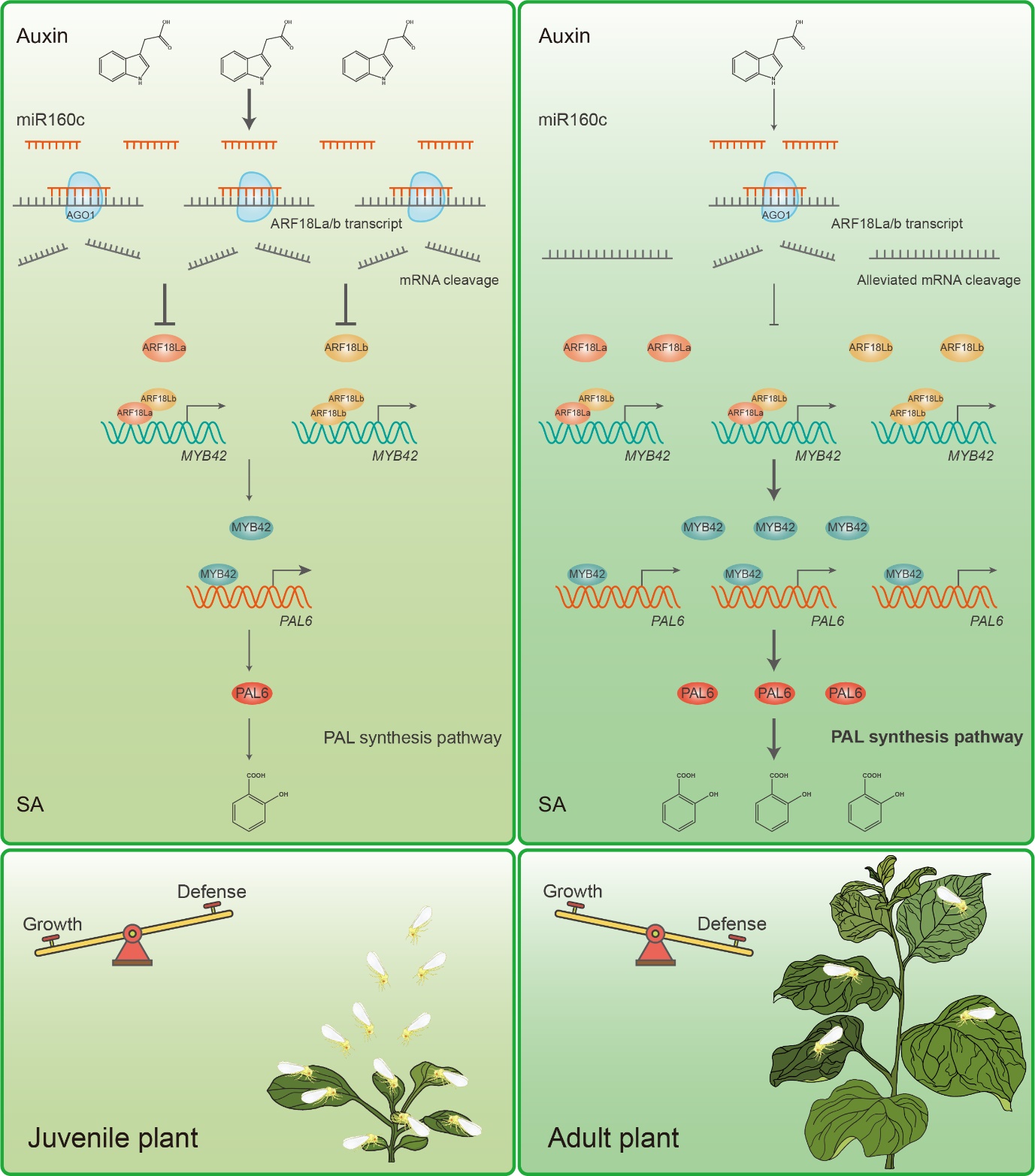

**Figure S11. Auxin-SA seesaw mediates the age-related balance between plant growth and herbivore defense**

At juvenile stages, plants exhibit high auxin levels, promoting the expression of *Nb*miR160c. This miRNA targets *NbARF18La/b* transcripts, leading to their degradation via the plant ARGONAUTE1 (AGO1) protein. As a result, NbARF18La/b is degraded at an early age, inhibiting downstream expression of *NbMYB42* and *NbPAL6*. This repression of the NbARF18La/b-NbMYB42-NbPAL6 module limits anti-whitefly SA accumulation in juvenile plants. As plants mature, the auxin levels decrease, lifting the repression of *NbARF18La/b*. These two interacting ARF TFs directly bind to the promoter of *NbMYB42*, activating its expression. NbMYB42 subsequently binds to the promoter of *NbPAL6*, enhancing its expression. NbPAL6 promotes SA biosynthesis, boosting SA-mediated defense against phloem-feeding insect whiteflies. The age-related rise in SA levels determines the ARR of plants. This model illustrates a seesaw-like mechanism for balancing growth and herbivore defense, with auxin dominance in early stages and SA-driven defense taking precedence as the plant ages.

**Supplementary Tables**

**Table S1. Basic information on identifying *NbPAL* genes.**

| Gene Name | Gene ID | CDS (bp) | Protein Length (aa) | Molecular Weight (Da) |
| --- | --- | --- | --- | --- |
| NbPAL1 | NbL01g24700.1 | 2040 | 679 | 73888.46 |
| NbPAL2 | NbL05g03880.1 | 2154 | 717 | 78005.01 |
| NbPAL3 | NbL10g04670.1 | 2154 | 717 | 78248.28 |
| NbPAL4 | NbL10g08440.1 | 2166 | 721 | 78734.94 |
| NbPAL5 | NbL12g19650.1 | 2154 | 717 | 78393.48 |
| NbPAL6 | NbL13g11330.1 | 2154 | 717 | 78233.31 |
| NbPAL7 | NbL14g10160.1 | 2139 | 712 | 77447.38 |
| NbPAL8 | NbL15g02170.1 | 2139 | 712 | 77438.39 |
| NbPAL9 | NbL15g24630.1 | 2154 | 717 | 78151.22 |

**Table S2. List of primers used in this study.**

| Primer name | Sequence (5’-3’) |
| --- | --- |
| q-mir160c-F | TGCCTGGCTCCCTGTATTTT |
| q-NbU6-F | GGAACGATACAGAGAAGATTAGCA |
| q-NbACTIN-F | CAATCCAGACACTGTACTTTCTCTC |
| q-NbACTIN-R | AAGCTGCAGGTATCCATGAGACTA |
| q-NbPR1-F | TGCAAATCAGAGAGCTGGGG |
| q-NbPR1-R | CACTTTTCCCGGTGCACAAG |
| q-NbICS1-F | CGTGCTTATGGGGCAATTCG |
| q-NbICS1-R | ATTGCCATCTGGTACGTGCA |
| q-NbICS2-F | CTCGCGAGTACTCATGTCCC |
| q-NbICS2-R | ATCAATGTCCGCAGCTGTCA |
| q-NbEPS1-F | GCCGGCGTAGAATTCACCTA |
| q-NbEPS1-R | GCCGACGTTATTGAGTGGGA |
| q-NbCBP60g-F | CGACCTTGGCAAGCAGTTTC |
| q-NbCBP60g-R | GGAGAAACTGGCTGGATGCT |
| q-NbSARD1-F | AGTTGGCCACTGTTGACACA |
| q-NbSARD1-R | AGTGTAGTTTGCTCCGGTGG |
| q-NbEDS5-F | CAAACACTGCTGCTGCACAT |
| q-NbEDS5-R | AGGGACTTCAGCAACATCCG |
| q-NbPBS3-F | AACCCCGAAATTACCGACCC |
| q-NbPBS3-R | AGCACCCGTGACAATTACGT |
| q-NbEPS1-F | GCCGGCGTAGAATTCACCTA |
| q-NbEPS1-R | GCCGACGTTATTGAGTGGGA |
| q-NbMYB42-F | GGTGAGAGCTCCTTGTTGTGA |
| q-NbMYB42-R | TCTCCAATTGCCATGGCCAT |
| q-NbPAL4-F | ATGCACTCAAGGGTAGCCAC |
| q-NbPAL4-R | CAGCTCCACCATCACCTCTG |
| q-NbPAL6-F | AGGAACAAGGCATTGCATGG |
| q-NbPAL6-R | TTTCCAATGGAGGCAAGTGC |
| q-NbPAL7-F | ACAATCAGGGACAAGGGCAG |
| q-NbPAL7-R | GACCTGTGAGTAAACCGGCA |
| q-NbPAL9-F | TGTTTACCTCTCCGTGGCAC |
| q-NbPAL9-R | CAGCATTAAGGGCCTCACCA |
| q-NbARF18La-R | TGATCCCATCTACTGGCCCA |
| q-NbARF18La-F | GAGACCAGTTCGACAAGCCA |
| q-NbARF18Lb-F | CAACTCGTGCTTTTCGGCAA |
| q-NbARF18Lb-R | AGTGCAGAACCTGAGCCATC |
| q-NbTAR2-F | GCTGAGACTGGAAACCGACA |
| q-NbTAR2-R | CGAATAAAATGGGGCAGCGG |
| q-NbYUC8-F | GCAAATGGCGCTGAAGTTGA |
| q-NbYUC8-R | TGCCAGAATTTCCACAGCCA |
| q-NbIAA29-F | CGTTTAAGGCTTGCGACGAG |
| q-NbIAA29-R | CCAATGGCCACTCCTTCCAT |
| pBI121-pNbPAL6-F | GACCATGATTACGCCAAGCTTGAATTCGTTCGGAAATTATGA |
| pBI121-pNbPAL6-R | ACCACCCGGGGATCCTCTAGAAAATGGAAAATATGAAAGAAAAAGA |
| pBI121-pNbARF18La-F | GACCATGATTACGCCAAGCTTCGCATGCAATCAAATATTATTACTCT |
| pBI121-pNbARF18La-R | ACCACCCGGGGATCCTCTAGATTGCCTTATCCAGTACACTTCT |
| pBI121-pNbARF18Lb-F | GACCATGATTACGCCAAGCTTGAGGTGAGGATTGCCCAAGC |
| pBI121-pNbARF18Lb-R | GGACTGACCACCCGGGGATCCCTTGATTTCAACAGAATTTTACAGA |
| pBI121-pNbMYB42-F | GCTATGACCATGATTACGCCAAGCTTTAGTGATTTGACTGTGCGTGA |
| pBI121-pNbMYB42-R | GACTGACCACCCGGGGATCCTCTAGATTTTGCTTTAATCACTTGCT |
| pBI121-pmir160-F | AGCTATGACCATGATTACGCCAAGCTTAATTTGACTATTACATAATTCCT |
| pBI121-pmir160-R | GGACTGACCACCCGGGGATCCCGAGGAAAATACGAAGTGAA |
| TRV-NbARF18La-F | GCTCTAGAACCCAGTTTCCAGCGAGTTG |
| TRV-NbARF18La-R | CGGGATCCTCTTGAAGACCTTTCTGCTCA |
| TRV-NbARF18Lb-F | GCTCTAGAAATCCAGCTGGTGGGAACTG |
| TRV-NbARF18Lb-R | CGGGATCCAAAAGGCATCTTGAACCTCATCC |
| TRV-NbPAL6-F | GCTCTAGATCTATGGTTCTATTTGAGGCT |
| TRV-NbPAL6-R | CGGGATCCGGCCCTAATGACCTCAATTTGA |
| TRV-NbMYB42-F | GAGTAAGGTTACCGAATTCTCTAGAAAAGACACTCCAAAAACAATCTTGA |
| TRV-NbMYB42-R | GTGAGCTCGGTACCGGATCCTTGGTCATCGATTGTAATAGTCGT |
| 1305-NbARF18La-F | TAAGTCCGGAGCTAGCTCTAGAATGGAGGAGGTTATGGAGAAATGT |
| 1305-NbARF18La-R | CGGTCCTCGAGACGTCTCTAGATGCAAACATGCTAAGTGGTCC |
| 1305-NbARF18Lb-F | TAAGTCCGGAGCTAGCTCTAGAATGATTACTTTTATGGATCCAAAGGAC |
| 1305-NbARF18Lb-R | CGGTCCTCGAGACGTCTCTAGACTCCCTGAGTCCAACATTATCG |
| 1305-NbPAL6-F | GCTCTAGAATGGAGTATGCAAATGGAGATTGT |
| 1305-NbPAL6-R | CGGGATCCGCAGAGTGGAAGAGGAGC |
| 1305-NbMYB42-F | GCTCTAGAATGGTGAGAGCTCCTTGTTGT |
| 1305-NbMYB42-R | CGGGATCCAAATTCCGGTAAATCTAGT |
| pABAi-pNbMYB42-F | AAAAAAAATGATGAATTGAAAAGCTTTAGTGATTTGACTGTGCGTGA |
| pABAi-pNbMYB42-R | TCGAGGTCGACAGATCCCCGGGTACCTTTTGCTTTAATCACTTGCT |
| pABAi-pNbPAL6-F | AAAAAAAATGATGAATTGAAAAGCTTGAATTCGTTCGGAAATTATGA |
| pABAi-pNbPAL6-R | TCGAGGTCGACAGATCCCCGGGTACCAAATGGAAAATATGAAAGAAAAAGA |
| pGAD-NbARF18La-F | CCGGGTGGGCATCGATACGGGATCCATGGAGGAGGTTATGGAGAAATGT |
| pGAD-NbARF18La-R | TATCTACGATTCATCTGCAGCTCGAGCTATGCAAACATGCTAAGTGGTCC |
| pGAD-NbARF18Lb-F | CCGGGTGGGCATCGATACGGGATCCATGATTACTTTTATGGATCCAAAGGAC |
| pGAD-NbARF18Lb-R | TATCTACGATTCATCTGCAGCTCGAGTTACTCCCTGAGTCCAACAT |
| pGAD-NbMYB42-F | CGGGATCCATGGTGAGAGCTCCTTGTTGT |
| pGAD-NbMYB42-R | CCCTCGAGTCAAAATTCCGGTAAATCTAGT |
| pGBK-NbARF18La-F | TGGCCATGGAGGCCGAATTCATGGAGGAGGTTATGGAGAAATGT |
| pGBK-NbARF18La-R | CGCTGCAGGTCGACGGATCCCTATGCAAACATGCTAAGTGGTCC |
| pGBK-NbARF18Lb-F | TGGCCATGGAGGCCGAATTCATGATTACTTTTATGGATCC |
| pGBK-NbARF18Lb-R | CGCTGCAGGTCGACGGATCCCTACTCCCTGAGTCCAACAT |
| pGreen0800-pNbPAL6-F | TCGAGGTCGACGGTATCGATAAGCTTGAATTCGTTCGGAAATTATGA |
| pGreen0800-pNbPAL6-R | GGCCGCTCTAGAACTAGTGGATCCAAATGGAAAATATGAAAGAAAAAGA |
| pGreen0800-pNbMYB42-F | TCGAGGTCGACGGTATCGATAAGCTTTAGTGATTTGACTGTGCGTGA |
| pGreen0800-pNbMYB42-R | GCGGCCGCTCTAGAACTAGTGGATCCTTTTGCTTTAATCACTTGCT |
| pGreen0800-pmir160-F | ACTCACTATAGGGCGAATTGGGTACCAATTTGACTATTACATAATTCCT |
| pGreen0800-pmir160-R | CGGCCGCTCTAGAACTAGTGGATCCCGAGGAAAATACGAAGTGAA |
| pET28a-SUMO-NbARF18La-F | GCTCACAGAGAACAGATTGGTGGAATGGAGGAGGTTATGGAGAAATGTG |
| pET28a-SUMO-NbARF18La-R | TGCCGCCACTACCACCACCACTACCACCGCCTTTTTCAAACTGCGGATGGCTCCATGCAAACATGCTAAGTGGTCCTGC |
| pMAL-NbARF18Lb-F | TCCAGGGGCCCCTGGGATCCATGATTACTTTTATGGATCC |
| pMAL-NbARF18Lb-R | CTGCCGCCACTACCACCACCACTACCACCGCCTTTTTCAAACTGCGGATGGCTCCACTCCCTGAGTCCAACATTATCGC |
| pGEX-NbMYB42-F | CTGGAAGTTCTGTTCCAGGGGCCCGGCTCCATGGTGAGAGCTCCTTGTTGTGA |
| pGEX-NbMYB42-R | CACTACCACCACCACTACCACCGCCTTTTTCAAACTGCGGATGGCTCCAAAATTCCGGTAAATCTAGTAAGTCCCCG |
| 1300-mir160-F | GGGGTACCCAGCAGCAGCCACAGCAAAA |
| 1300-mir160-R | CGGGATCCGCTGCTGATGCTGATGCCAT |
| 1300-mimic160-F | GGTACCTGGCCATCCCCTA |
| 1300-mimic160-R | GGATCCCGGAAGCAAATTTACA |
| Bifc-NbARF18La-F | CCATTTACGAACGATAGTTAATTAACATGGAGGAGGTTATGGAGAAATGT |
| Bifc-NbARF18La-R | CCACCACTGCCACCTCCTCCACTAGTTGCAAACATGCTAAGTGGTCC |
| Bifc-NbARF18Lb-F | CCATTTACGAACGATAGTTAATTAACATGATTACTTTTATGGATCCAAAGGAC |
| Bifc-NbARF18Lb-R | CCACCACTGCCACCTCCTCCACTAGTCTCCCTGAGTCCAACATTATCG |
| 35S-GFP-mir160-F | AGGACCGGTCCCGGGGGATCCATGGTGAGCAAGGGCGAG |
| GFP-NbARF18La-site-R | GGGCGGCCGCTTTAAGATCTATGCCTGGCTCTCTGTATGC |
| GFP-NbARF18La-mutant site-R | GGGCGGCCGCTTTAAGATCTATGTCTCGCGCTCTCTAGGC |
| GFP-NbARF18Lb-site-R | GGGCGGCCGCTTTAAGATCTATGCCTGGCTCCCTGCAT |
| GFP-NbARF18Lb-mutant site-R | GGGCGGCCGCTTTAAGATCTATGTCTTGCGCCTTGAATGC |
| 35S-GUS-mir160-F | GGAGAGAACACGGGGGACTCTAGAATGTTACGTCCTGTAGAAACCCC |
| GUS-NbARF18La-site-R | AACGATCGGGGAAATTCGAGCTCTCAATGCCTGGCTCTCTGTATGCC |
| GUS-NbARF18La-mutant site-R | AACGATCGGGGAAATTCGAGCTCTCAATGTCTCGCGCTCTCTAGGCC |
| GUS-NbARF18Lb-site-R | AACGATCGGGGAAATTCGAGCTCTCAATGCCTGGCTCCCTGCATGCC |
| GUS-NbARF18Lb-mutant site-R | AACGATCGGGGAAATTCGAGCTCTCAATGTCTTGCGCCTTGAATGCC |
| 35S-LUC-mir160-F | CATGCCATGGAAGACGCCAA |
| LUC-NbARF18La-site-R | GCTCTAGAATGCCTGGCTCTCTGTATGCCATTACACGGCGATCTTTCCG |
| LUC-NbARF18La-mutant site-R | GCTCTAGAATGTCTCGCGCTCTCTAGGCCATTACACGGCGATCTTTCCG |
| LUC-NbARF18Lb-site-R | GCTCTAGAATGCCTGGCTCCCTGCATGCCATTACACGGCGATCTTTCCG |
| LUC-NbARF18Lb-mutant site-R | GCTCTAGAATGTCTTGCGCCTTGAATGCCATTACACGGCGATCTTTCCG |
| ChIP-pNbMYB42-P1-F | AGTGATTTGACTGTGCGTGA |
| ChIP-pNbMYB42-P1-R | AAAACGGCACAAATGGTCCG |
| ChIP-pNbMYB42-P2-F | AACTTGAGAACCCACGACCC |
| ChIP-pNbMYB42-P2-R | CCTGCATGGAAATTCATGAAATCT |
| ChIP-pNbMYB42-P3-F | ACTCCTTCAGATTTTGACGTGA |
| ChIP-pNbMYB42-P3-R | ACCTTATTTACGAGGATTACACTGAGT |
| ChIP-pNbPAL6-P1-F | TCGGTCTCAACGCGTTAATGA |
| ChIP-pNbPAL6-P1-R | ACTTTCCCTTCACCAACATGA |
| ChIP-pNbPAL6-P2-F | ATGACGTGGTGCTGTCGTTG |
| ChIP-pNbPAL6-P2-R | TTTCCCTCGTAAAATACCGCA |
| ChIP-pNbPAL6-P3-F | CGCCAGTAAGTTATTATCGCGC |
| ChIP-pNbPAL6-P3-R | CGTTGGATTGCACATGGAGA |
| ChIP-pNbPAL6-P4-F | TCTCCATGTGCAATCCAACG |
| ChIP-pNbPAL6-P4-R | TGTTGAGCCAAAATGTGAGTGT |
| ChIP-pNbPAL6-P5-F | TGGCTCAACAAACTCAGGAAATT |
| ChIP-pNbPAL6-P5-R | AATGTGTGTTGCGAACTTCTC |
| EMSA-pPAL6-F | GTGAGGGTTGAAGGGTGGTTGTTGAGAGGAAGCCCGCTTACTTTTTTTGG |
| EMSA-pPAL6-R | GGTTTTTTTCATTCGCCCGAAGGAGAGTTGTTGGTGGGAAGTTGGGAGTG |
| EMSA-pPAL6-mF | GTGAGGGTTGAAGAAAAAAAAAAAAAAAAAAGCCCGCTTACTTTTTTTGG |
| EMSA-pPAL6-mR | GGTTTTTTTCATTCGCCCGAAAAAAAAAAAAAAAAAAGAAGTTGGGAGTG |
| EMSA-pMYB42-F | GGACAATTTTTCTTTATTTTCTTTTTTGTCTCCAGTAAAATAACG |
| EMSA-pMYB42-R | GCAATAAAATGACCTCTGTTTTTTCTTTTATTTCTTTTTAACAGG |
| EMSA-pMYB42-mF | GGACAATTTTTCTTTATTTTCTTTTTAAAAAACAGTAAAATAACG |
| EMSA-pMYB42-mR | GCAATAAAATGACAAAAAATTTTTCTTTTATTTCTTTTTAACAGG |
